# Genomic determinants of Furin cleavage in diverse European SARS-related bat coronaviruses

**DOI:** 10.1101/2021.12.15.472779

**Authors:** Anna-Lena Sander, Andres Moreira-Soto, Stoian Yordanov, Ivan Toplak, Andrea Balboni, Ramón Seage Ameneiros, Victor Corman, Christian Drosten, Jan Felix Drexler

## Abstract

The furin cleavage site in SARS-CoV-2 is unique within the *Severe acute respiratory syndrome–related coronavirus* (*SrC*) species. We re-assessed diverse *SrC* from European horseshoe bats and reveal molecular determinants such as purine richness, RNA secondary structures and viral quasispecies potentially enabling furin cleavage. Furin cleavage thus likely emerged from the *SrC* bat reservoir via molecular mechanisms conserved across reservoir-bound RNA viruses, supporting a natural origin of SARS-CoV-2.

---

Emerging coronaviruses of recent or regular zoonotic origin include the betacoronaviruses Severe acute respiratory syndrome coronavirus (SARS-CoV), Middle East respiratory syndrome coronavirus (MERS-CoV) and Severe acute respiratory syndrome coronavirus 2 (SARS-CoV-2) (1). SARS-CoV-2 is unique in its high transmissibility (2). SARS-CoV and SARS-CoV-2 belong to the species *Severe acute respiratory syndrome–related coronavirus* (*SrC*), subgenus Sarbecovirus, and both use angiotensin-converting enzyme 2 (ACE2) as main cellular receptor (3). In contrast to SARS-CoV, only SARS-CoV-2 contains a functional polybasic furin cleavage site (FCS) between the two subunits of the viral spike glycoprotein (4). The existence of a FCS has led to various hypotheses regarding the evolution of SARS-CoV-2, including conjectures about the possibility of a non-natural origin from laboratory experiments (5, 6). Furin cleavage is thought to be essential for entry into human lung cells and may also determine the efficiency of infection of the upper respiratory tract and consequent transmissibility of SARS-CoV-2 (7). So far, SARS-CoV-2 is unique among *SrC*, as even its closest known relatives, the bat coronavirus RaTG13 and the pangolin coronaviruses, lack a FCS (8). The natural hosts of *SrC* are horseshoe bats, widely distributed in the Old World (9). We and others previously showed that *SrC* in European horseshoe bats are conspecific but distinct from those detected in Asia (10–13). Here, we describe the S1/S2 genomic region encompassing the FCS in SARS-CoV-2 in ten unique European bat-associated *SrC* in comparison to other sarbecoviruses and mammalian coronaviruses.

We re-accessed stored original fecal samples from four horseshoe bat species (*Rhinolophus hipposideros*, *R. euryale*, *R. ferrumequinum*, *R. blasii*) collected in Italy, Bulgaria, Spain, and Slovenia during 2008-2009 and amplified an 816 nucleotide fragment of the viral *RNA-dependent RNA polymerase (RdRp)* of ten unique coronaviruses previously described (12, 13). Taxonomic classification based on this fragment showed that all ten coronaviruses belonged to the species *SrC* (**Supplementary Table 1**). In a representative phylogeny that covered the diversity of known *SrC*, based on a partial S2-genomic region encompassing 495 nucleotides, European bat-associated *SrC* formed a sister clade to Chinese bat-associated *SrC* (**Figure panel A, Supplementary Figure 1**). Sequence comparison of the S1/S2 genomic region revealed remnants of a polybasic FCS motif (R-X-X-R) at the S1/S2 boundary in 12 of 71 unique bat-associated *SrC* from Europe, Asia, and Africa with higher genetic diversity in European than in Asian bat-associated *SrC* (**Figure panel B**).

**Figure.**
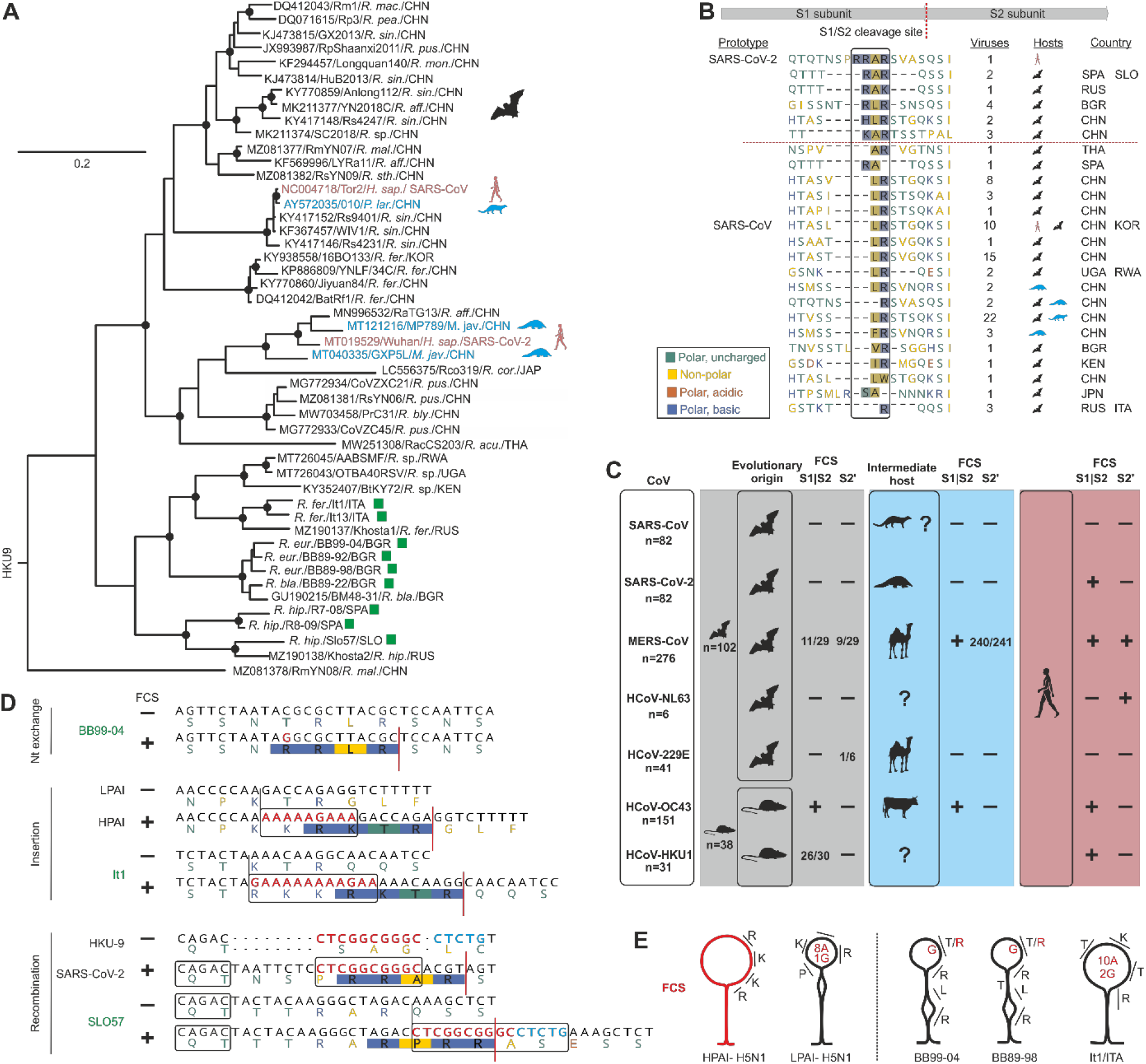
Evolutionary relationships of SARS-related and other mammalian coronaviruses. **A**) Approximately-Maximum-likelihood (ML) phylogeny based on a partial S2-fragment (495bp) of representative sequences of the complete diversity of *SrC* shown in the phylogeny in the supplementary figure. European horseshoe bat *SrC* generated within this study are shown with green squares and without Accession numbers, SARS-CoV and SARS-CoV-2 are given in red, and *SrC* from civets and pangolins in blue type. Sequences are named as followed: GenBank Acc. number/strain name/host species/country of detection. Circles at nodes indicate support of grouping in ≥ 90% of 1,000 bootstrap replicates. Scale bar represents nucleotide substitutions per site. **B**) Bat-associated *SrC* harbor remnants of a FCS between the spike subunits S1 and S2. Genomic regions at the interface of the S1 and S2 domains and the S2’ position of the spike protein of 91 unique *SrC* were aligned using Mafft (23). A scheme of the spike protein and its subunits S1 and S2 shows the position of the FCS in SARS-CoV-2. Conserved amino acids of the FCS are highlighted within the box. Red line separates sequences with and without remnants of FCS. **C**) FCS conservation at the S1/S2 interface and the S2’ site in human coronaviruses and closely related animal coronaviruses. FCS predictions are shown in the Supplementary Table. If not all tested sequences of one host category were predicted to contain a FCS, counts are given. **D**) Potential generation of FCS in different European bat *SrC* by nt exchange, insertion due to external stem loop structures as shown for HPAI (19) or recombination with HKU9, as speculated for SARS-CoV-2 (22). FCS scores were 0.65 for BB99-04, 0.69 for IT1 and 0.68 for SLO57. **E**) Predicted RNA secondary structures of the polybasic cleavage site regions of AIV and *SrC* that acquire FCS through nucleotide substitutions or insertions. Amino acids corresponding to codons forming the FCS are shown. Nucleotides and corresponding amino acids that change to a FCS through nucleotide substitution or insertions are marked in red. *H. sap*., *Homo sapiens*; *M. jav*., *Manis javanica*; *P. lar*., *Paguma larvata*; *R. acu*., *Rhinolophus acuminatus*; *R. aff*., *Rhinolophus affinus*; *R. bla*., *Rhinolophus blasii*; *R. bly*., *Rhinolophus blythi*; *R. cor*., *Rhinolophus cornutus*; *R. eur*., *Rhinolophus euryale*; *R. fer*., *Rhinolophus ferrumequinum*; *R. hip*., *Rhinolophus hipposideros*; *R. mac*., *Rhinolophus macrotis*; *R. mal*., *Rhinolophus malayanus*; *R. mon*., *Rhinolophus monoceros*; *R. pea*., *Rhinolophus pearsonii*; *R. pus*., *Rhinolophus pusillus*; *R. sin*., *Rhinolophus sinicus*; *R.* sp., *Rhinolophus* species*; R. sth*., *Rhinolophus stheno*; BGR, Bulgaria; CHN, China; ITA, Italy; JPN, Japan; KEN, Kenya; KOR, Korea; RUS, Russia; RWA, Ruanda; SLO, Slovenia; SPA, Spain; THA, Thailand; UGA, Uganda.

Next to the S1/S2 FCS, the coronaviral spike protein can also be activated by host cell proteases at the N-terminal S2 (S2’) genomic region (8). Only MERS-CoV contains a FCS at both the S1/S2 and the S2’ sites (14), whereas the other coronaviruses contain an intact FCS at either the S1/S2 or S2’ site. To better understand the evolution of FCS at both the S1/S2 and S2’ sites within human coronaviruses we investigated the genomic regions encompassing potential FCS within human-associated coronaviruses and related viruses found in their ancestral and intermediate hosts. Within bat-associated CoVs, only 10/102 (10%) and 11/102 (11%) showed a FCS in either the S2’ or the S1/S2 genomic region, respectively, suggesting circulation of a broad genetic diversity in the genomic region encoding potential FCS in the mammalian coronavirus bat reservoir (**Figure panel C** and **Supplementary Table 2**). For example, the human coronaviruses 229E and NL63 differed in their S2′ FCS integrity from viruses that represent (a sample of) the ancestral viral diversity in bats. Interestingly, a FCS at the S1/S2 boundary is not present in studied bat CoVs with the exception of some bat-associated MERS-related CoVs (15), suggesting that a FCS may not provide a fitness advantage in some or most bat hosts. A hypothetical turnover of FCS in the animal reservoir and fixation of FCS after host switches is reminiscent of the prototypic example of an RNA virus gaining pathogenicity via acquisition of a FCS, the avian Influenza A virus (IAV). IAVs are distinguished into low-pathogenic avian influenza virus (LPAI) and high-pathogenic avian influenza virus (HPAI). HPAIs are defined by the existence of a polybasic FCS at the hemagglutinin (HA) cleavage site. LPAI evolve into HPAI within the reservoir or the new host (16, 17) by acquisition of a FCS through three different molecular mechanisms: recombination with cellular or other RNA molecules, multiple nucleotide insertions, or nucleotide substitutions (17–19). Both insertions and nucleotide substitutions in the HA of IAVs are facilitated by a stem-loop secondary RNA structure enclosing the FCS and a high adenine/guanine content in the external loop structure (19). Importantly, genomic surrogates of all three mechanisms present in IAV are given in European bat-associated *SrC* (**Figure panel D**). First, predicted RNA secondary structures among some European bat-associated *SrC* and HPAI sequences suggest similarities in the genomic determinants of FCS acquisition between avian HPAI and bat *SrC* (**Figure panel E** and **Supplementary Figure 2**). Additionally, a single non-synonymous nucleotide substitution (an A to G transversion) in the external loop of the RNA secondary structure would suffice to allow furin cleavage in two European bat-associated *SrC* (termed BB99-04 and BB89-98) (**Figure panel E**). Deep sequencing of this genome position revealed presence of this transversion in 0.004% and 0.006% of the total reads in those two viruses (**Table 1**). On the one hand, single nucleotide variants affording the emergence of furin cleavage in LPAI were found at a comparatively low frequency (0.0028%), which may imply the potential for emergence of FCS in those European bat *SrC* strains (19). On the other hand, the occurrence of those transversions was within the error rate of viral polymerases used for cDNA synthesis and amplification (20). Whether bat *SrC* quasispecies harboring a functional FCS exist within European bat hosts thus requires careful examination. Next, a more than 60% adenine/guanine content in the stem loop structure of another European *SrC* (termed It1) may facilitate an insertion of an adenine/guanine stretch and thus the acquisition of a FCS (**Figure panel D and E**), comparable to the insertion leading to HPAI outbreaks in the US in 2016/2017 (17, 21). Finally, as speculated for the origin of the S1/S2 FCS of SARS-CoV-2 (22), recombination with other bat coronaviruses such as HKU9 would result in the acquisition of a S1/S2 FCS. The existence of the palindromic sequence CAGAC in another European *SrC* (termed SLO57) comparable to SARS-CoV-2 (**Figure panel D**) suggests that this genomic region may serve as an RNA signal for a recombination breakpoint in some European bat *SrC* (22).

**Table 1.** Single nucleotide variants within two European *SrC* at spike position Thr670.

| BB99-04 | Consensus sequence | A | C | G |
| --- | --- | --- | --- | --- |
|  | A (%) | 146506 (99.801) | 8 (0.005) | 159 (0.108) |
|  | T (%) | 19 (0.013) | 187 (0.127) | 13 (0.009) |
|  | C (%) | 7 (0.005) | 147250 (99.858) | 10 (0.007) |
|  | G (%) | 259 (0.176) | **6\* (0.004)** | 146817 (99.872) |
|  | Total reads | 146798 | 147459 | 147005 |
| BB89-98 | Consensus sequence | A | C | A |
|  | A (%) | 117032 (99.593) | 1 (0.001) | 116893 (99.745) |
|  | T (%) | 47 (0.0400) | 38 (0.032) | 13 (0.011) |
|  | C (%) | 10 (0.009) | 117540 (99.961) | 8 (0.007) |
|  | G (%) | 421 (0.358) | **7\*\* (0.006)** | 278 (0.237) |
|  | Total reads | 117510 | 117586 | 117192 |

#One of the reads was not paired-end
\*This single nucleotide variant leads to a non-synonymous exchange generating a FCS motif in this virus

In sum, our analyses support the acquisition of FCS in European bat-associated *SrC* via mechanisms similar to the genesis of HPAI in their avian reservoir. This supports a natural evolutionary origin of SARS-CoV-2 in bats with or without the involvement of intermediary hosts. Ecological studies into bats are likely to identify other sarbecoviruses harboring functional FCS. The zoonotic potential of such sarbecoviruses deserves investigation to identify variants potentially posing threats to human health.

## Acknowledgments

This study was funded by the German Research Foundation (DFG) (DR 810/7-1, DR 772/12-1), as well as the German Ministry of Research (01KI1723A). We thank Mara Battilani of the department of Veterinary Medical Sciences from University of Bologna, Italy, Danijela Černe of the Institute of Microbiology and Parasitology from University of Ljubljana, Slovenia, Florian Gloza-Rausch of the Behavioral Ecology and Bioacostics Lab at Museum für Naturkunde, Berlin, Germany for providing sample material and Monika Eschbach-Bludau for technical assistance.

## Methods

### Sample collection and processing

Bats were sampled with mist nets using minimally invasive methods under appropriate permits as described earlier (10, 11, 13). Specimens were screened for the presence of viral RNA of the genus Coronavirus by using reverse-transcription-PCR (RT-PCR) as described previously (24), amplifying 455 bp fragments of the *RNA-dependent RNA polymerase* (RdRp) gene. For further phylogenetic analyses these amplicons were extended to an 816 bp fragment towards the 5’ end (12). Nucleotide sequences were deposited in GenBank with accession numbers KC633198, KC633201-205, KC633209, KC633212 and KC633217.

S1/S2 genomic regions were characterized using a hemi-nested RT-PCR assay flanking the S1/S2 and S2’ site (690 nt; pos 23,442-24,112 in SARS-CoV-2 Wuhan strain Acc. Number MT019529) using the following oligonucleotides: panSARS-S1S2-F1 (TDGCTGTTGTHTAYCARGATGT), panSARS-S1S2-F2 (CARGATGTWAAYTGYACWGATGT) and panSARS-S1S2-R (AGDCCATTRAACTTYTGHGCACA). Briefly, RNA was reverse transcribed for 30 min at 50°C using the SSIII One-Step Kit (Thermo Fisher) followed by 45 PCR cycles of 94°C for 15 seconds, 58°C for 45 seconds and 72°C for 1 minute. The 2nd round PCR was performed at the same conditions as the 1st round without reverse transcription. PCR amplicons were Sanger sequenced.

To detect single nucleotide variants within the S1/S2 site, PCR amplicons were sequenced using the Illumina NovaSeq 6000 Sequencing System with the NovaSeq 6000 SP Reagent Kit (500 cycles). Sequence reads obtained from the library were mapped against their corresponding S1/S2 genomic sequence obtained after PCR amplification in Geneious 9.1.8.

Nucleotide sequences obtained within this study were deposited in GenBank with accession numbers XXX to XXX.

### Phylogenetic analyses

A tblastx search of the complete spike sequence of the Bulgarian *SrC* (GenBank Acc No. GU190215) within the Taxa ID 11118 (*Coronaviridae*) excluding the Taxa ID 2697049 (SARS-CoV-2) was performed on 21 June 2021. Hits with percentage identities below 80% were non-*SrC* sequences and were thus not included to the dataset. SARS-CoV sequences, experimentally infected or clones as well as sequences with less than 27,000 nt or gaps in the spike protein were excluded, resulting in 80 sequences. One reference sequence of each SARS-CoV and SARS-CoV-2 as well as the nine sequences from European bats generated within this study were additionally added, resulting in a final dataset of 91 sequences. Because Coronaviruses frequently recombine (25) only the S2 region (495 nt) of the 690 nt amplified fragment was used for the phylogenetic analyses. Maximum-likelihood phylogenies were generated using FastTree Version 2.1.10 using a GTR substitution model and 1,000 bootstrap replicates. Local support values are based on the Shimodaira-Hasegawa (SH) test (26).

### In silico analyses

Secondary structures were modeled using the UNAfold web server (27). Furin cleavage sites were predicted using ProP v.1.0b ProPeptide Cleavage Site Prediction (28). Sequences were retrieved from the NCBI Taxonomy website downloading all sequences of the species Duvinacovirus (HCoV-229E), Merbecovirus (MERS-CoV), Setracovirus (HCoV-NL63) and Embecovirus (HCoV-HKU1 and HCoV-OC43). Duplicates, non-complete genomes, sequences from animal experiences and clones were excluded from all datasets. For Sarbecoviruses, the dataset of the tBlastx search used for the *SrC*-phylogeny of the Supplementary Figure 1 was used. Accession numbers of viruses tested in ProP are listed in the Supplementary Material.

## References

1. Jo WK, Drosten C, Drexler JF. The evolutionary dynamics of endemic human coronaviruses. Virus Evol. 2021 Jan;7(1):veab020.

2. Jones TC, Biele G, Muhlemann B, Veith T, Schneider J, Beheim-Schwarzbach J, et al. Estimating infectiousness throughout SARS-CoV-2 infection course. Science. 2021 May 25.

3. Hoffmann M, Kleine-Weber H, Schroeder S, Kruger N, Herrler T, Erichsen S, et al. SARS-CoV-2 Cell Entry Depends on ACE2 and TMPRSS2 and Is Blocked by a Clinically Proven Protease Inhibitor. Cell. 2020 Mar 4.

4. Coutard B, Valle C, de Lamballerie X, Canard B, Seidah NG, Decroly E. The spike glycoprotein of the new coronavirus 2019-nCoV contains a furin-like cleavage site absent in CoV of the same clade. Antiviral Res. 2020 Feb 10;176:104742.

5. Calisher CH, Carroll D, Colwell R, Corley RB, Daszak P, Drosten C, et al. Science, not speculation, is essential to determine how SARS-CoV-2 reached humans. Lancet. 2021 Jul 17;398(10296):209–11.

6. Bloom JD, Chan YA, Baric RS, Bjorkman PJ, Cobey S, Deverman BE, et al. Investigate the origins of COVID-19. Science. 2021 May 14;372(6543):694.

7. Hoffmann M, Kleine-Weber H, Pohlmann S. A Multibasic Cleavage Site in the Spike Protein of SARS-CoV-2 Is Essential for Infection of Human Lung Cells. Mol Cell. 2020 May 21;78(4):779–84 e5.

8. Wu Y, Zhao S. Furin cleavage sites naturally occur in coronaviruses. Stem Cell Res. 2020 Dec 9;50:102115.

9. Jo WK, de Oliveira-Filho EF, Rasche A, Greenwood AD, Osterrieder K, Drexler JF. Potential zoonotic sources of SARS-CoV-2 infections. Transbound Emerg Dis. 2020 Oct 9.

10. Balboni A, Palladini A, Bogliani G, Battilani M. Detection of a virus related to betacoronaviruses in Italian greater horseshoe bats. Epidemiology and infection. 2011 Feb;139(2):216–9.

11. Rihtaric D, Hostnik P, Steyer A, Grom J, Toplak I. Identification of SARS-like coronaviruses in horseshoe bats (Rhinolophus hipposideros) in Slovenia. Arch Virol. 2010 Apr;155(4):507–14.

12. Drexler JF, Gloza-Rausch F, Glende J, Corman VM, Muth D, Goettsche M, et al. Genomic characterization of severe acute respiratory syndrome-related coronavirus in European bats and classification of coronaviruses based on partial RNA-dependent RNA polymerase gene sequences. J Virol. 2010 Nov;84(21):11336–49.

13. Muth D, Corman VM, Roth H, Binger T, Dijkman R, Gottula LT, et al. Attenuation of replication by a 29 nucleotide deletion in SARS-coronavirus acquired during the early stages of human-to-human transmission. Sci Rep. 2018 Oct 11;8(1):15177.

14. Millet JK, Whittaker GR. Host cell entry of Middle East respiratory syndrome coronavirus after two-step, furin-mediated activation of the spike protein. Proc Natl Acad Sci U S A. 2014 Oct 21;111(42):15214–9.

15. Stout AE, Millet JK, Stanhope MJ, Whittaker GR. Furin cleavage sites in the spike proteins of bat and rodent coronaviruses: Implications for virus evolution and zoonotic transfer from rodent species. One Health. 2021 Dec;13:100282.

16. Monne I, Fusaro A, Nelson MI, Bonfanti L, Mulatti P, Hughes J, et al. Emergence of a highly pathogenic avian influenza virus from a low-pathogenic progenitor. J Virol. 2014 Apr;88(8):4375–88.

17. Lee DH, Torchetti MK, Killian ML, Swayne DE. Deep sequencing of H7N8 avian influenza viruses from surveillance zone supports H7N8 high pathogenicity avian influenza was limited to a single outbreak farm in Indiana during 2016. Virology. 2017 Jul;507:216–9.

18. Suarez DL, Senne DA, Banks J, Brown IH, Essen SC, Lee CW, et al. Recombination resulting in virulence shift in avian influenza outbreak, Chile. Emerg Infect Dis. 2004 Apr;10(4):693–9.

19. Nao N, Yamagishi J, Miyamoto H, Igarashi M, Manzoor R, Ohnuma A, et al. Genetic Predisposition To Acquire a Polybasic Cleavage Site for Highly Pathogenic Avian Influenza Virus Hemagglutinin. mBio. 2017 Feb 14;8(1).

20. Orton RJ, Wright CF, Morelli MJ, King DJ, Paton DJ, King DP, et al. Distinguishing low frequency mutations from RT-PCR and sequence errors in viral deep sequencing data. BMC Genomics. 2015 Mar 24;16:229.

21. Lee DH, Torchetti MK, Killian ML, Berhane Y, Swayne DE. Highly Pathogenic Avian Influenza A(H7N9) Virus, Tennessee, USA, March 2017. Emerg Infect Dis. 2017 Nov;23(11).

22. Gallaher WR. A palindromic RNA sequence as a common breakpoint contributor to copy-choice recombination in SARS-COV-2. Arch Virol. 2020 Oct;165(10):2341–8.

23. Katoh K, Misawa K, Kuma K, Miyata T. MAFFT: a novel method for rapid multiple sequence alignment based on fast Fourier transform. Nucleic acids research. 2002 Jul 15;30(14):3059–66.

24. de Souza Luna LK, Heiser V, Regamey N, Panning M, Drexler JF, Mulangu S, et al. Generic detection of coronaviruses and differentiation at the prototype strain level by reverse transcription-PCR and nonfluorescent low-density microarray. J Clin Microbiol. 2007 Mar;45(3):1049–52.

25. Vakulenko Y, Deviatkin A, Drexler JF, Lukashev A. Modular Evolution of Coronavirus Genomes. Viruses. 2021;13(7):1270.

26. Shimodaira H, Hasegawa M. Multiple Comparisons of Log-Likelihoods with Applications to Phylogenetic Inference. Molecular Biology and Evolution. 1999;16(8):1114–.

27. Zuker M. Mfold web server for nucleic acid folding and hybridization prediction. Nucleic acids research. 2003 Jul 1;31(13):3406–15.

28. Duckert P, Brunak S, Blom N. Prediction of proprotein convertase cleavage sites. Protein Eng Des Sel. 2004 Jan;17(1):107–12.

