## Supplementary information for "Genomic determinants of Furin cleavage in diverse European SARS-related bat coronaviruses"

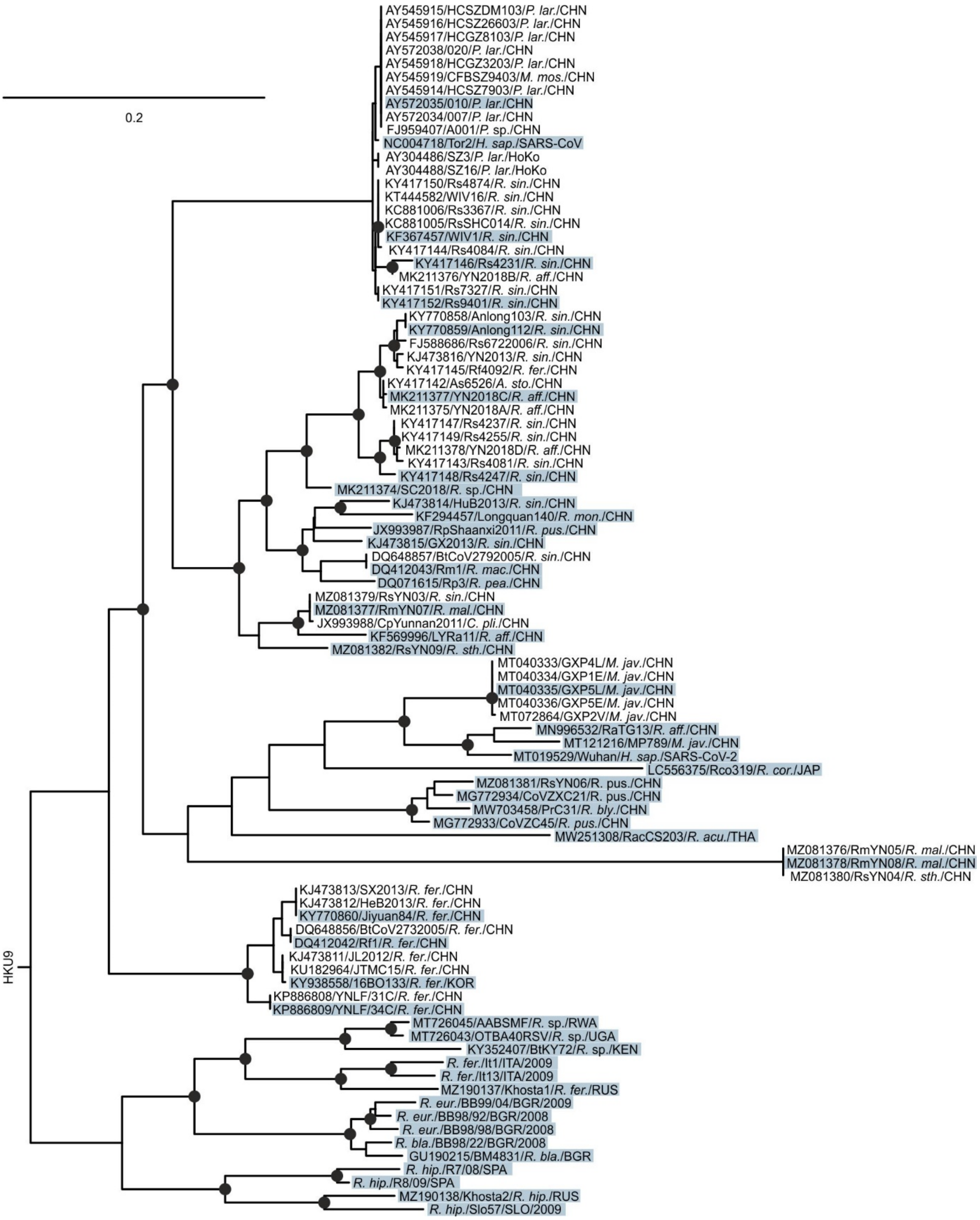


**Supplementary Figure 1.** ML phylogeny showing the complete diversity of members of the species *SARS-related Coronavirus* (*SrC*). The final dataset comprised 91 sequences (2 human, 12 civet, 6

pangolin and 71 bat-associated *SrC*). Sequences are named as followed: GenBank Acc. number/strain name/host species/country of detection. Circles at nodes indicate support of grouping in ≥ 90% of 1,000 bootstrap replicates. Scale bar represents nucleotide substitutions per site. Sequences highlighted in blue were chosen for phylogenetic analysis shown in Figure panel A, representing the complete diversity of *SrC*. *A. sto.*, *Aselliscus stoliczkanus; C. pli.*, *Chaerephon plicatus; H. sap.*, *Homo sapiens*; *M. jav.*, *Manis javanica*; *M. mos*, *Melogale moschata*; *P. lar.*, *Paguma larvata*; *P.* sp.; *Paguma* species; *R. acu.*, *Rhinolophus acuminatus*; *R. aff.*, *Rhinolophus affinus*; *R. bla.*, *Rhinolophus blasii*; *R. bly.*, *Rhinolophus blythi*; *R. cor.*, *Rhinolophus cornutus*; *R. eur.*, *Rhinolophus euryale*; *R. fer.*, *Rhinolophus ferrumequinum*; *R. hip.*, *Rhinolophus hipposideros*; *R. mac.*, *Rhinolophus macrotis*; *R. mal.*, *Rhinolophus malayanus*; *R. mon.*, *Rhinolophus monoceros*; *R. pea.*, *Rhinolophus pearsonii*; *R. pus.*, *Rhinolophus pusillus*; *R. sin.*, *Rhinolophus sinicus*; *R.* sp., *Rhinolophus* species*; R. sth.*, *Rhinolophus stheno*; BGR, Bulgaria; CHN, China; ITA, Italy; JPN, Japan; KEN, Kenya; KOR, Korea; RUS, Russia; RWA, Ruanda; SLO, Slovenia; SPA, Spain; THA, Thailand; UGA, Uganda.

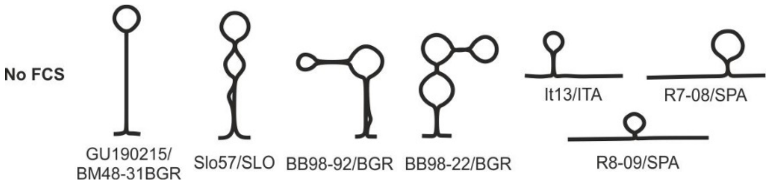


**Supplementary Figure 2.** Predicted RNA secondary structures of the polybasic cleavage site regions of *SrC* that are not predicted to acquire FCS through nucleotide substitutions or insertions.

**Supplementary table 1. Amino acid identities of a partial *RdRp* fragment of European bat coronaviruses and prototype betacoronaviruses for species delineation.**

| % amino acid sequence identity (range): | | | |
|---|---|---|---|
| Within European bat CoVs | European bat CoVs compared to | | |
| | SARS-CoV (AY274119) | SARS-CoV-2 (MT019529) | MERS-CoV (NC019843) |
| 97.8-100% | 97.4-99.3 | 96.7-97.8 | 76.4-76.5 |

**Supplementary table 2.** FCS prediction in human coronaviruses and their ancestors

| Related HCoV | FCS region | Accession number | Prototype | Host category | Order | FCS | ProP score |
|---|---|---|---|---|---|---|---|
| 229E | S2 | KT253272 | | Evolutionary origin | Chiroptera | GSGRTGR\|SV | 0.675 |
| HKU1 | S1S2 | AC000192 | | Evolutionary origin | Rodentia | KSRRARR\|SV | 0.884 |
| HKU1 | S1S2 | FJ647219 | | Evolutionary origin | Rodentia | KSRRARR\|SV | 0.884 |
| HKU1 | S1S2 | FJ647220 | | Evolutionary origin | Rodentia | KSRRARR\|SV | 0.884 |
| HKU1 | S1S2 | FJ647221 | | Evolutionary origin | Rodentia | KSRRARR\|SV | 0.884 |
| HKU1 | S1S2 | FJ647224 | | Evolutionary origin | Rodentia | KSRRARR\|SV | 0.884 |
| HKU1 | S1S2 | FJ647226 | | Evolutionary origin | Rodentia | KSRRARR\|SV | 0.884 |

| | | | | | | | |
|---|---|---|---|---|---|---|---|
| HKU1 | S1S2 | JX169866 | | Evolutionary origin | Rodentia | KSRRARR\|SV | 0.884 |
| HKU1 | S1S2 | JX169867 | | Evolutionary origin | Rodentia | KSRRARR\|SV | 0.884 |
| HKU1 | S1S2 | MW620427 | | Evolutionary origin | Rodentia | KSRRARR\|SV | 0.884 |
| HKU1 | S1S2 | FJ938068 | | Evolutionary origin | Rodentia | TAHRARR\|SV | 0.881 |
| HKU1 | S1S2 | JF792617 | | Evolutionary origin | Rodentia | TAHRARR\|SV | 0.881 |
| HKU1 | S1S2 | NC012936 | | Evolutionary origin | Rodentia | TAHRARR\|SV | 0.881 |
| HKU1 | S1S2 | AB551247 | | Evolutionary origin | Rodentia | KARRARR\|SV | 0.875 |
| HKU1 | S1S2 | MF416379 | | Evolutionary origin | Rodentia | XAHRARR\|SV | 0.871 |
| HKU1 | S1S2 | JF792616 | | Evolutionary origin | Rodentia | IAHRARR\|SV | 0.867 |
| HKU1 | S1S2 | KF850449 | | Evolutionary origin | Rodentia | IAHRARR\|SV | 0.867 |
| HKU1 | S1S2 | FJ647223 | | Evolutionary origin | Rodentia | TSHRARR\|SI | 0.865 |
| HKU1 | S1S2 | MW773844 | | Evolutionary origin | Rodentia | TSHRARR\|SI | 0.865 |
| HKU1 | S1S2 | FJ647218 | | Evolutionary origin | Rodentia | KSRRAHR\|SV | 0.808 |
| HKU1 | S1S2 | FJ647222 | | Evolutionary origin | Rodentia | KSRRAHR\|SV | 0.808 |
| HKU1 | S1S2 | FJ647225 | | Evolutionary origin | Rodentia | KSRRAHR\|SV | 0.808 |
| HKU1 | S1S2 | FJ647227 | | Evolutionary origin | Rodentia | KSRRAHR\|SV | 0.808 |
| HKU1 | S1S2 | MF618252 | | Evolutionary origin | Rodentia | KSRRAHR\|SV | 0.808 |
| HKU1 | S1S2 | FJ884687 | | Evolutionary origin | Rodentia | KSRRADR\|SV | 0.798 |
| HKU1 | S1S2 | NC001846 | | Evolutionary origin | Rodentia | KSRRADR\|SV | 0.798 |
| HKU1 | S1S2 | NC048217 | | Evolutionary origin | Rodentia | KSRRAHR\|SV | 0.808 |
| HKU1 | S1S2 | NC006577 | HCoV-HKU1 | | Primates | SSRRKRR\|SI | 0.877 |
| HKU1 | S1S2 | NC006577 | HCoV-HKU1 | | Primates | SSSRRKR\|RS | 0.654 |
| MERS | S1S2 | MH002341 | | Evolutionary origin | Chiroptera | TSSRVRR\|AT | 0.853 |
| MERS | S1S2 | EF065509 | | Evolutionary origin | Chiroptera | TSSRVRR\|AT | 0.822 |
| MERS | S1S2 | NC_009020 | | Evolutionary origin | Chiroptera | TSSRVRR\|AT | 0.822 |
| MERS | S1S2 | KJ473820 | | Evolutionary origin | Chiroptera | TSSRLRR\|AT | 0.811 |
| MERS | S1S2 | MH002340 | | Evolutionary origin | Chiroptera | TPSRVLR\|AA | 0.746 |
| MERS | S1S2 | MH002342 | | Evolutionary origin | Chiroptera | PSARLAR\|SA | 0.701 |
| MERS | S1S2 | EF065510 | | Evolutionary origin | Chiroptera | TSTRFRR\|AT | 0.588 |
| MERS | S1S2 | EF065511 | | Evolutionary origin | Chiroptera | TSTRFRR\|AT | 0.588 |
| MERS | S1S2 | EF065512 | | Evolutionary origin | Chiroptera | TSTRFRR\|AT | 0.588 |
| MERS | S1S2 | KC869678 | | Evolutionary origin | Chiroptera | TNLRSGR\|ST | 0.572 |
| MERS | S1S2 | MF593268 | | Evolutionary origin | Chiroptera | TNLRSGR\|ST | 0.572 |
| MERS | S1S2 | MH002342 | | Evolutionary origin | Chiroptera | RLARSAR\|SG | 0.509 |
| MERS | S1S2 | MK967708 | | Intermediate host | Artiodactyla | LTPRSVR\|SV | 0.607 |
| MERS | S1S2 | KJ477103 | | Intermediate host | Artiodactyla | LTPRSVR\|SV | 0.563 |
| MERS | S1S2 | KU740200 | | Intermediate host | Artiodactyla | LTPRSVR\|SV | 0.563 |
| MERS | S1S2 | KY581695 | | Intermediate host | Artiodactyla | LTPRSVR\|SV | 0.563 |
| MERS | S1S2 | KY581696 | | Intermediate host | Artiodactyla | LTPRSVR\|SV | 0.563 |
| MERS | S1S2 | KY581697 | | Intermediate host | Artiodactyla | LTPRSVR\|SV | 0.563 |
| MERS | S1S2 | KY581698 | | Intermediate host | Artiodactyla | LTPRSVR\|SV | 0.563 |
| MERS | S1S2 | KY581699 | | Intermediate host | Artiodactyla | LTPRSVR\|SV | 0.563 |
| MERS | S1S2 | KY581700 | | Intermediate host | Artiodactyla | LTPRSVR\|SV | 0.563 |
| MERS | S1S2 | KY673149 | | Intermediate host | Artiodactyla | LTPRSVR\|SV | 0.563 |
| MERS | S1S2 | MF598594 | | Intermediate host | Artiodactyla | LTPRSVR\|SV | 0.563 |
| MERS | S1S2 | MF598595 | | Intermediate host | Artiodactyla | LTPRSVR\|SV | 0.563 |
| MERS | S1S2 | MF598596 | | Intermediate host | Artiodactyla | LTPRSVR\|SV | 0.563 |
| MERS | S1S2 | MF598597 | | Intermediate host | Artiodactyla | LTPRSVR\|SV | 0.563 |
| MERS | S1S2 | MF598598 | | Intermediate host | Artiodactyla | LTPRSVR\|SV | 0.563 |
| MERS | S1S2 | MF598599 | | Intermediate host | Artiodactyla | LTPRSVR\|SV | 0.563 |
| MERS | S1S2 | MF598600 | | Intermediate host | Artiodactyla | LTPRSVR\|SV | 0.563 |
| MERS | S1S2 | MF598601 | | Intermediate host | Artiodactyla | LTPRSVR\|SV | 0.563 |
| MERS | S1S2 | MF598602 | | Intermediate host | Artiodactyla | LTPRSVR\|SV | 0.563 |
| MERS | S1S2 | MF598603 | | Intermediate host | Artiodactyla | LTPRSVR\|SV | 0.563 |
| MERS | S1S2 | MF598604 | | Intermediate host | Artiodactyla | LTPRSVR\|SV | 0.563 |
| MERS | S1S2 | MF598605 | | Intermediate host | Artiodactyla | LTPRSVR\|SV | 0.563 |
| MERS | S1S2 | MF598606 | | Intermediate host | Artiodactyla | LTPRSVR\|SV | 0.563 |
| MERS | S1S2 | MF598607 | | Intermediate host | Artiodactyla | LTPRSVR\|SV | 0.563 |
| MERS | S1S2 | MF598608 | | Intermediate host | Artiodactyla | LTPRSVR\|SV | 0.563 |
| MERS | S1S2 | MF598609 | | Intermediate host | Artiodactyla | LTPRSVR\|SV | 0.563 |
| MERS | S1S2 | MF598610 | | Intermediate host | Artiodactyla | LTPRSVR\|SV | 0.563 |
| MERS | S1S2 | MF598611 | | Intermediate host | Artiodactyla | LTPRSVR\|SV | 0.563 |
| MERS | S1S2 | MF598612 | | Intermediate host | Artiodactyla | LTPRSVR\|SV | 0.563 |
| MERS | S1S2 | MF598613 | | Intermediate host | Artiodactyla | LTPRSVR\|SV | 0.563 |
| MERS | S1S2 | MF598614 | | Intermediate host | Artiodactyla | LTPRSVR\|SV | 0.563 |
| MERS | S1S2 | MF598615 | | Intermediate host | Artiodactyla | LTPRSVR\|SV | 0.563 |
| MERS | S1S2 | MF598616 | | Intermediate host | Artiodactyla | LTPRSVR\|SV | 0.563 |
| MERS | S1S2 | MF598617 | | Intermediate host | Artiodactyla | LTPRSVR\|SV | 0.563 |
| MERS | S1S2 | MF598618 | | Intermediate host | Artiodactyla | LTPRSVR\|SV | 0.563 |
| MERS | S1S2 | MF598619 | | Intermediate host | Artiodactyla | LTPRSVR\|SV | 0.563 |
| MERS | S1S2 | MF598620 | | Intermediate host | Artiodactyla | LTPRSVR\|SV | 0.563 |
| MERS | S1S2 | MF598621 | | Intermediate host | Artiodactyla | LTPRSVR\|SV | 0.563 |
| MERS | S1S2 | MF598622 | | Intermediate host | Artiodactyla | LTPRSVR\|SV | 0.563 |

| | | | | | | | |
|---|---|---|---|---|---|---|---|
| MERS | S1S2 | MF598623 | | Intermediate host | Artiodactyla | LTPRSVR\|SV | 0.563 |
| MERS | S1S2 | MF598624 | | Intermediate host | Artiodactyla | LTPRSVR\|SV | 0.563 |
| MERS | S1S2 | MF598625 | | Intermediate host | Artiodactyla | LTPRSVR\|SV | 0.563 |
| MERS | S1S2 | MF598626 | | Intermediate host | Artiodactyla | LTPRSVR\|SV | 0.563 |
| MERS | S1S2 | MF598627 | | Intermediate host | Artiodactyla | LTPRSVR\|SV | 0.563 |
| MERS | S1S2 | MF598628 | | Intermediate host | Artiodactyla | LTPRSVR\|SV | 0.563 |
| MERS | S1S2 | MF598629 | | Intermediate host | Artiodactyla | LTPRSVR\|SV | 0.563 |
| MERS | S1S2 | MF598630 | | Intermediate host | Artiodactyla | LTPRSVR\|SV | 0.563 |
| MERS | S1S2 | MF598631 | | Intermediate host | Artiodactyla | LTPRSVR\|SV | 0.563 |
| MERS | S1S2 | MF598632 | | Intermediate host | Artiodactyla | LTPRSVR\|SV | 0.563 |
| MERS | S1S2 | MF598633 | | Intermediate host | Artiodactyla | LTPRSVR\|SV | 0.563 |
| MERS | S1S2 | MF598634 | | Intermediate host | Artiodactyla | LTPRSVR\|SV | 0.563 |
| MERS | S1S2 | MF598635 | | Intermediate host | Artiodactyla | LTPRSVR\|SV | 0.563 |
| MERS | S1S2 | MF598636 | | Intermediate host | Artiodactyla | LTPRSVR\|SV | 0.563 |
| MERS | S1S2 | MF598637 | | Intermediate host | Artiodactyla | LTPRSVR\|SV | 0.563 |
| MERS | S1S2 | MF598638 | | Intermediate host | Artiodactyla | LTPRSVR\|SV | 0.563 |
| MERS | S1S2 | MF598639 | | Intermediate host | Artiodactyla | LTPRSVR\|SV | 0.563 |
| MERS | S1S2 | MF598640 | | Intermediate host | Artiodactyla | LTPRSVR\|SV | 0.563 |
| MERS | S1S2 | MF598641 | | Intermediate host | Artiodactyla | LTPRSVR\|SV | 0.563 |
| MERS | S1S2 | MF598642 | | Intermediate host | Artiodactyla | LTPRSVR\|SV | 0.563 |
| MERS | S1S2 | MF598643 | | Intermediate host | Artiodactyla | LTPRSVR\|SV | 0.563 |
| MERS | S1S2 | MF598644 | | Intermediate host | Artiodactyla | LTPRSVR\|SV | 0.563 |
| MERS | S1S2 | MF598645 | | Intermediate host | Artiodactyla | LTPRSVR\|SV | 0.563 |
| MERS | S1S2 | MF598646 | | Intermediate host | Artiodactyla | LTPRSVR\|SV | 0.563 |
| MERS | S1S2 | MF598647 | | Intermediate host | Artiodactyla | LTPRSVR\|SV | 0.563 |
| MERS | S1S2 | MF598648 | | Intermediate host | Artiodactyla | LTPRSVR\|SV | 0.563 |
| MERS | S1S2 | MF598649 | | Intermediate host | Artiodactyla | LTPRSVR\|SV | 0.563 |
| MERS | S1S2 | MF598650 | | Intermediate host | Artiodactyla | LTPRSVR\|SV | 0.563 |
| MERS | S1S2 | MF598651 | | Intermediate host | Artiodactyla | LTPRSVR\|SV | 0.563 |
| MERS | S1S2 | MF598652 | | Intermediate host | Artiodactyla | LTPRSVR\|SV | 0.563 |
| MERS | S1S2 | MF598653 | | Intermediate host | Artiodactyla | LTPRSVR\|SV | 0.563 |
| MERS | S1S2 | MF598654 | | Intermediate host | Artiodactyla | LTPRSVR\|SV | 0.563 |
| MERS | S1S2 | MF598655 | | Intermediate host | Artiodactyla | LTPRSVR\|SV | 0.563 |
| MERS | S1S2 | MF598656 | | Intermediate host | Artiodactyla | LTPRSVR\|SV | 0.563 |
| MERS | S1S2 | MF598657 | | Intermediate host | Artiodactyla | LTPRSVR\|SV | 0.563 |
| MERS | S1S2 | MF598658 | | Intermediate host | Artiodactyla | LTPRSVR\|SV | 0.563 |
| MERS | S1S2 | MF598659 | | Intermediate host | Artiodactyla | LTPRSVR\|SV | 0.563 |
| MERS | S1S2 | MF598660 | | Intermediate host | Artiodactyla | LTPRSVR\|SV | 0.563 |
| MERS | S1S2 | MF598661 | | Intermediate host | Artiodactyla | LTPRSVR\|SV | 0.563 |
| MERS | S1S2 | MF598662 | | Intermediate host | Artiodactyla | LTPRSVR\|SV | 0.563 |
| MERS | S1S2 | MF598663 | | Intermediate host | Artiodactyla | LTPRSVR\|SV | 0.563 |
| MERS | S1S2 | MF598664 | | Intermediate host | Artiodactyla | LTPRSVR\|SV | 0.563 |
| MERS | S1S2 | MF598665 | | Intermediate host | Artiodactyla | LTPRSVR\|SV | 0.563 |
| MERS | S1S2 | MF598666 | | Intermediate host | Artiodactyla | LTPRSVR\|SV | 0.563 |
| MERS | S1S2 | MF598667 | | Intermediate host | Artiodactyla | LTPRSVR\|SV | 0.563 |
| MERS | S1S2 | MF598668 | | Intermediate host | Artiodactyla | LTPRSVR\|SV | 0.563 |
| MERS | S1S2 | MF598669 | | Intermediate host | Artiodactyla | LTPRSVR\|SV | 0.563 |
| MERS | S1S2 | MF598670 | | Intermediate host | Artiodactyla | LTPRSVR\|SV | 0.563 |
| MERS | S1S2 | MF598671 | | Intermediate host | Artiodactyla | LTPRSVR\|SV | 0.563 |
| MERS | S1S2 | MF598672 | | Intermediate host | Artiodactyla | LTPRSVR\|SV | 0.563 |
| MERS | S1S2 | MF598673 | | Intermediate host | Artiodactyla | LTPRSVR\|SV | 0.563 |
| MERS | S1S2 | MF598674 | | Intermediate host | Artiodactyla | LTPRSVR\|SV | 0.563 |
| MERS | S1S2 | MF598675 | | Intermediate host | Artiodactyla | LTPRSVR\|SV | 0.563 |
| MERS | S1S2 | MF598676 | | Intermediate host | Artiodactyla | LTPRSVR\|SV | 0.563 |
| MERS | S1S2 | MF598677 | | Intermediate host | Artiodactyla | LTPRSVR\|SV | 0.563 |
| MERS | S1S2 | MF598678 | | Intermediate host | Artiodactyla | LTPRSVR\|SV | 0.563 |
| MERS | S1S2 | MF598679 | | Intermediate host | Artiodactyla | LTPRSVR\|SV | 0.563 |
| MERS | S1S2 | MF598680 | | Intermediate host | Artiodactyla | LTPRSVR\|SV | 0.563 |
| MERS | S1S2 | MF598681 | | Intermediate host | Artiodactyla | LTPRSVR\|SV | 0.563 |
| MERS | S1S2 | MF598682 | | Intermediate host | Artiodactyla | LTPRSVR\|SV | 0.563 |
| MERS | S1S2 | MF598683 | | Intermediate host | Artiodactyla | LTPRSVR\|SV | 0.563 |
| MERS | S1S2 | MF598684 | | Intermediate host | Artiodactyla | LTPRSVR\|SV | 0.563 |
| MERS | S1S2 | MF598685 | | Intermediate host | Artiodactyla | LTPRSVR\|SV | 0.563 |
| MERS | S1S2 | MF598686 | | Intermediate host | Artiodactyla | LTPRSVR\|SV | 0.563 |
| MERS | S1S2 | MF598687 | | Intermediate host | Artiodactyla | LTPRSVR\|SV | 0.563 |
| MERS | S1S2 | MF598688 | | Intermediate host | Artiodactyla | LTPRSVR\|SV | 0.563 |
| MERS | S1S2 | MF598689 | | Intermediate host | Artiodactyla | LTPRSVR\|SV | 0.563 |
| MERS | S1S2 | MF598690 | | Intermediate host | Artiodactyla | LTPRSVR\|SV | 0.563 |
| MERS | S1S2 | MF598691 | | Intermediate host | Artiodactyla | LTPRSVR\|SV | 0.563 |
| MERS | S1S2 | MF598692 | | Intermediate host | Artiodactyla | LTPRSVR\|SV | 0.563 |
| MERS | S1S2 | MF598693 | | Intermediate host | Artiodactyla | LTPRSVR\|SV | 0.563 |
| MERS | S1S2 | MF598694 | | Intermediate host | Artiodactyla | LTPRSVR\|SV | 0.563 |
| MERS | S1S2 | MF598695 | | Intermediate host | Artiodactyla | LTPRSVR\|SV | 0.563 |

| | | | | | | | |
|---|---|---|---|---|---|---|---|
| MERS | S1S2 | MF598696 | | Intermediate host | Artiodactyla | LTPRSVR\|SV | 0.563 |
| MERS | S1S2 | MF598697 | | Intermediate host | Artiodactyla | LTPRSVR\|SV | 0.563 |
| MERS | S1S2 | MF598698 | | Intermediate host | Artiodactyla | LTPRSVR\|SV | 0.563 |
| MERS | S1S2 | MF598699 | | Intermediate host | Artiodactyla | LTPRSVR\|SV | 0.563 |
| MERS | S1S2 | MF598700 | | Intermediate host | Artiodactyla | LTPRSVR\|SV | 0.563 |
| MERS | S1S2 | MF598701 | | Intermediate host | Artiodactyla | LTPRSVR\|SV | 0.563 |
| MERS | S1S2 | MF598702 | | Intermediate host | Artiodactyla | LTPRSVR\|SV | 0.563 |
| MERS | S1S2 | MF598703 | | Intermediate host | Artiodactyla | LTPRSVR\|SV | 0.563 |
| MERS | S1S2 | MF598704 | | Intermediate host | Artiodactyla | LTPRSVR\|SV | 0.563 |
| MERS | S1S2 | MF598705 | | Intermediate host | Artiodactyla | LTPRSVR\|SV | 0.563 |
| MERS | S1S2 | MF598706 | | Intermediate host | Artiodactyla | LTPRSVR\|SV | 0.563 |
| MERS | S1S2 | MF598707 | | Intermediate host | Artiodactyla | LTPRSVR\|SV | 0.563 |
| MERS | S1S2 | MF598708 | | Intermediate host | Artiodactyla | LTPRSVR\|SV | 0.563 |
| MERS | S1S2 | MF598709 | | Intermediate host | Artiodactyla | LTPRSVR\|SV | 0.563 |
| MERS | S1S2 | MF598710 | | Intermediate host | Artiodactyla | LTPRSVR\|SV | 0.563 |
| MERS | S1S2 | MF598711 | | Intermediate host | Artiodactyla | LTPRSVR\|SV | 0.563 |
| MERS | S1S2 | MF598712 | | Intermediate host | Artiodactyla | LTPRSVR\|SV | 0.563 |
| MERS | S1S2 | MF598713 | | Intermediate host | Artiodactyla | LTPRSVR\|SV | 0.563 |
| MERS | S1S2 | MF598714 | | Intermediate host | Artiodactyla | LTPRSVR\|SV | 0.563 |
| MERS | S1S2 | MF598715 | | Intermediate host | Artiodactyla | LTPRSVR\|SV | 0.563 |
| MERS | S1S2 | MF598716 | | Intermediate host | Artiodactyla | LTPRSVR\|SV | 0.563 |
| MERS | S1S2 | MF598717 | | Intermediate host | Artiodactyla | LTPRSVR\|SV | 0.563 |
| MERS | S1S2 | MF598718 | | Intermediate host | Artiodactyla | LTPRSVR\|SV | 0.563 |
| MERS | S1S2 | MF598719 | | Intermediate host | Artiodactyla | LTPRSVR\|SV | 0.563 |
| MERS | S1S2 | MF598720 | | Intermediate host | Artiodactyla | LTPRSVR\|SV | 0.563 |
| MERS | S1S2 | MF598721 | | Intermediate host | Artiodactyla | LTPRSVR\|SV | 0.563 |
| MERS | S1S2 | MF598722 | | Intermediate host | Artiodactyla | LTPRSVR\|SV | 0.563 |
| MERS | S1S2 | MG923469 | | Intermediate host | Artiodactyla | LTPRSVR\|SV | 0.563 |
| MERS | S1S2 | MG923470 | | Intermediate host | Artiodactyla | LTPRSVR\|SV | 0.563 |
| MERS | S1S2 | MG923471 | | Intermediate host | Artiodactyla | LTPRSVR\|SV | 0.563 |
| MERS | S1S2 | MG923472 | | Intermediate host | Artiodactyla | LTPRSVR\|SV | 0.563 |
| MERS | S1S2 | MG923473 | | Intermediate host | Artiodactyla | LTPRSVR\|SV | 0.563 |
| MERS | S1S2 | MG923474 | | Intermediate host | Artiodactyla | LTPRSVR\|SV | 0.563 |
| MERS | S1S2 | MG923475 | | Intermediate host | Artiodactyla | LTPRSVR\|SV | 0.563 |
| MERS | S1S2 | MG923476 | | Intermediate host | Artiodactyla | LTPRSVR\|SV | 0.563 |
| MERS | S1S2 | MG923477 | | Intermediate host | Artiodactyla | LTPRSVR\|SV | 0.563 |
| MERS | S1S2 | MG923478 | | Intermediate host | Artiodactyla | LTPRSVR\|SV | 0.563 |
| MERS | S1S2 | MG923479 | | Intermediate host | Artiodactyla | LTPRSVR\|SV | 0.563 |
| MERS | S1S2 | MG923480 | | Intermediate host | Artiodactyla | LTPRSVR\|SV | 0.563 |
| MERS | S1S2 | MG923481 | | Intermediate host | Artiodactyla | LTPRSVR\|SV | 0.563 |
| MERS | S1S2 | MH259485 | | Intermediate host | Artiodactyla | LTPRSVR\|SV | 0.563 |
| MERS | S1S2 | MH259486 | | Intermediate host | Artiodactyla | LTPRSVR\|SV | 0.563 |
| MERS | S1S2 | MH734114 | | Intermediate host | Artiodactyla | LTPRSVR\|SV | 0.563 |
| MERS | S1S2 | MH734115 | | Intermediate host | Artiodactyla | LTPRSVR\|SV | 0.563 |
| MERS | S1S2 | MK357908 | | Intermediate host | Artiodactyla | LTPRSVR\|SV | 0.563 |
| MERS | S1S2 | MK357909 | | Intermediate host | Artiodactyla | LTPRSVR\|SV | 0.563 |
| MERS | S1S2 | MK564474 | | Intermediate host | Artiodactyla | LTPRSVR\|SV | 0.563 |
| MERS | S1S2 | MK564475 | | Intermediate host | Artiodactyla | LTPRSVR\|SV | 0.563 |
| MERS | S1S2 | MN654970 | | Intermediate host | Artiodactyla | LTPRSVR\|SV | 0.563 |
| MERS | S1S2 | MN654971 | | Intermediate host | Artiodactyla | LTPRSVR\|SV | 0.563 |
| MERS | S1S2 | MN654972 | | Intermediate host | Artiodactyla | LTPRSVR\|SV | 0.563 |
| MERS | S1S2 | MN654973 | | Intermediate host | Artiodactyla | LTPRSVR\|SV | 0.563 |
| MERS | S1S2 | MN654974 | | Intermediate host | Artiodactyla | LTPRSVR\|SV | 0.563 |
| MERS | S1S2 | MN654975 | | Intermediate host | Artiodactyla | LTPRSVR\|SV | 0.563 |
| MERS | S1S2 | MN654976 | | Intermediate host | Artiodactyla | LTPRSVR\|SV | 0.563 |
| MERS | S1S2 | MN654977 | | Intermediate host | Artiodactyla | LTPRSVR\|SV | 0.563 |
| MERS | S1S2 | MN654978 | | Intermediate host | Artiodactyla | LTPRSVR\|SV | 0.563 |
| MERS | S1S2 | MN654979 | | Intermediate host | Artiodactyla | LTPRSVR\|SV | 0.563 |
| MERS | S1S2 | MN654980 | | Intermediate host | Artiodactyla | LTPRSVR\|SV | 0.563 |
| MERS | S1S2 | MN654981 | | Intermediate host | Artiodactyla | LTPRSVR\|SV | 0.563 |
| MERS | S1S2 | MN654982 | | Intermediate host | Artiodactyla | LTPRSVR\|SV | 0.563 |
| MERS | S1S2 | MN654983 | | Intermediate host | Artiodactyla | LTPRSVR\|SV | 0.563 |
| MERS | S1S2 | MN654984 | | Intermediate host | Artiodactyla | LTPRSVR\|SV | 0.563 |
| MERS | S1S2 | MN654985 | | Intermediate host | Artiodactyla | LTPRSVR\|SV | 0.563 |
| MERS | S1S2 | MN654986 | | Intermediate host | Artiodactyla | LTPRSVR\|SV | 0.563 |
| MERS | S1S2 | MN654987 | | Intermediate host | Artiodactyla | LTPRSVR\|SV | 0.563 |
| MERS | S1S2 | MN654988 | | Intermediate host | Artiodactyla | LTPRSVR\|SV | 0.563 |
| MERS | S1S2 | MN654989 | | Intermediate host | Artiodactyla | LTPRSVR\|SV | 0.563 |
| MERS | S1S2 | MN654990 | | Intermediate host | Artiodactyla | LTPRSVR\|SV | 0.563 |
| MERS | S1S2 | MN654991 | | Intermediate host | Artiodactyla | LTPRSVR\|SV | 0.563 |
| MERS | S1S2 | MN654992 | | Intermediate host | Artiodactyla | LTPRSVR\|SV | 0.563 |
| MERS | S1S2 | MN654993 | | Intermediate host | Artiodactyla | LTPRSVR\|SV | 0.563 |
| MERS | S1S2 | MN654994 | | Intermediate host | Artiodactyla | LTPRSVR\|SV | 0.563 |

| | | | | | | | |
|---|---|---|---|---|---|---|---|
| MERS | S1S2 | MN654996 | | Intermediate host | Artiodactyla | LTPRSVR\|SV | 0.563 |
| MERS | S1S2 | MN654997 | | Intermediate host | Artiodactyla | LTPRSVR\|SV | 0.563 |
| MERS | S1S2 | MN654998 | | Intermediate host | Artiodactyla | LTPRSVR\|SV | 0.563 |
| MERS | S1S2 | MN654999 | | Intermediate host | Artiodactyla | LTPRSVR\|SV | 0.563 |
| MERS | S1S2 | MN655000 | | Intermediate host | Artiodactyla | LTPRSVR\|SV | 0.563 |
| MERS | S1S2 | MN655001 | | Intermediate host | Artiodactyla | LTPRSVR\|SV | 0.563 |
| MERS | S1S2 | MN655002 | | Intermediate host | Artiodactyla | LTPRSVR\|SV | 0.563 |
| MERS | S1S2 | MN655003 | | Intermediate host | Artiodactyla | LTPRSVR\|SV | 0.563 |
| MERS | S1S2 | MN655004 | | Intermediate host | Artiodactyla | LTPRSVR\|SV | 0.563 |
| MERS | S1S2 | MN655005 | | Intermediate host | Artiodactyla | LTPRSVR\|SV | 0.563 |
| MERS | S1S2 | MN655007 | | Intermediate host | Artiodactyla | LTPRSVR\|SV | 0.563 |
| MERS | S1S2 | MN655008 | | Intermediate host | Artiodactyla | LTPRSVR\|SV | 0.563 |
| MERS | S1S2 | MN655009 | | Intermediate host | Artiodactyla | LTPRSVR\|SV | 0.563 |
| MERS | S1S2 | MN655010 | | Intermediate host | Artiodactyla | LTPRSVR\|SV | 0.563 |
| MERS | S1S2 | MN655011 | | Intermediate host | Artiodactyla | LTPRSVR\|SV | 0.563 |
| MERS | S1S2 | MN655012 | | Intermediate host | Artiodactyla | LTPRSVR\|SV | 0.563 |
| MERS | S1S2 | MN655013 | | Intermediate host | Artiodactyla | LTPRSVR\|SV | 0.563 |
| MERS | S1S2 | MN655014 | | Intermediate host | Artiodactyla | LTPRSVR\|SV | 0.563 |
| MERS | S1S2 | MN655015 | | Intermediate host | Artiodactyla | LTPRSVR\|SV | 0.563 |
| MERS | S1S2 | MN655016 | | Intermediate host | Artiodactyla | LTPRSVR\|SV | 0.563 |
| MERS | S1S2 | MN655017 | | Intermediate host | Artiodactyla | LTPRSVR\|SV | 0.563 |
| MERS | S1S2 | MN758606 | | Intermediate host | Artiodactyla | LTPRSVR\|SV | 0.563 |
| MERS | S1S2 | MN758607 | | Intermediate host | Artiodactyla | LTPRSVR\|SV | 0.563 |
| MERS | S1S2 | MN758608 | | Intermediate host | Artiodactyla | LTPRSVR\|SV | 0.563 |
| MERS | S1S2 | MN758609 | | Intermediate host | Artiodactyla | LTPRSVR\|SV | 0.563 |
| MERS | S1S2 | MN758610 | | Intermediate host | Artiodactyla | LTPRSVR\|SV | 0.563 |
| MERS | S1S2 | MN758611 | | Intermediate host | Artiodactyla | LTPRSVR\|SV | 0.563 |
| MERS | S1S2 | MN758612 | | Intermediate host | Artiodactyla | LTPRSVR\|SV | 0.563 |
| MERS | S1S2 | MN758613 | | Intermediate host | Artiodactyla | LTPRSVR\|SV | 0.563 |
| MERS | S1S2 | MN758614 | | Intermediate host | Artiodactyla | LTPRSVR\|SV | 0.563 |
| MERS | S1S2 | MT226600 | | Intermediate host | Artiodactyla | LTPRSVR\|SV | 0.563 |
| MERS | S1S2 | MT226601 | | Intermediate host | Artiodactyla | LTPRSVR\|SV | 0.563 |
| MERS | S1S2 | MT226602 | | Intermediate host | Artiodactyla | LTPRSVR\|SV | 0.563 |
| MERS | S1S2 | MT226603 | | Intermediate host | Artiodactyla | LTPRSVR\|SV | 0.563 |
| MERS | S1S2 | MT226604 | | Intermediate host | Artiodactyla | LTPRSVR\|SV | 0.563 |
| MERS | S1S2 | MT226605 | | Intermediate host | Artiodactyla | LTPRSVR\|SV | 0.563 |
| MERS | S1S2 | MT226606 | | Intermediate host | Artiodactyla | LTPRSVR\|SV | 0.563 |
| MERS | S1S2 | MT226607 | | Intermediate host | Artiodactyla | LTPRSVR\|SV | 0.563 |
| MERS | S1S2 | MW086527 | | Intermediate host | Artiodactyla | LTPRSVR\|SV | 0.563 |
| MERS | S1S2 | MW086528 | | Intermediate host | Artiodactyla | LTPRSVR\|SV | 0.563 |
| MERS | S1S2 | MW086529 | | Intermediate host | Artiodactyla | LTPRSVR\|SV | 0.563 |
| MERS | S1S2 | MW086530 | | Intermediate host | Artiodactyla | LTPRSVR\|SV | 0.563 |
| MERS | S1S2 | MW086531 | | Intermediate host | Artiodactyla | LTPRSVR\|SV | 0.563 |
| MERS | S1S2 | MW086532 | | Intermediate host | Artiodactyla | LTPRSVR\|SV | 0.563 |
| MERS | S1S2 | MW086533 | | Intermediate host | Artiodactyla | LTPRSVR\|SV | 0.563 |
| MERS | S1S2 | MW086534 | | Intermediate host | Artiodactyla | LTPRSVR\|SV | 0.563 |
| MERS | S1S2 | MW086535 | | Intermediate host | Artiodactyla | LTPRSVR\|SV | 0.563 |
| MERS | S1S2 | MW086538 | | Intermediate host | Artiodactyla | LTPRSVR\|SV | 0.563 |
| MERS | S1S2 | MW086539 | | Intermediate host | Artiodactyla | LTPRSVR\|SV | 0.563 |
| MERS | S1S2 | MW545527 | | Intermediate host | Artiodactyla | LTPRSVR\|SV | 0.563 |
| MERS | S1S2 | MW545528 | | Intermediate host | Artiodactyla | LTPRSVR\|SV | 0.563 |
| MERS | S1S2 | MG923466 | | Intermediate host | Artiodactyla | FTPRSVR\|SV | 0.536 |
| MERS | S1S2 | MG923467 | | Intermediate host | Artiodactyla | FTPRSVR\|SV | 0.536 |
| MERS | S1S2 | MN654995 | | Intermediate host | Artiodactyla | FTPRSVR\|SV | 0.536 |
| MERS | S1S2 | MN655006 | | Intermediate host | Artiodactyla | FTPRSVR\|SV | 0.536 |
| MERS | S1S2 | MG923468 | | Intermediate host | Artiodactyla | FTPRSVR\|SV | 0.522 |
| MERS | S1S2 | MH371127 | MERS-CoV | | Primates | LTPRSVR\|SV | 0.563 |
| MERS | S1S2 | NC019843 | MERS-CoV | | Primates | LTPRSVR\|SV | 0.563 |
| MERS | S1S2 | NC038294 | MERS-CoV | | Primates | LTPRSVR\|SV | 0.563 |
| MERS | S2 | MH002340 | | Evolutioanry origin | Chiroptera | TGERKYR\|SA | 0.575 |
| MERS | S2 | KJ473820 | | Evolutioanry origin | Chiroptera | TGERKYR\|SA | 0.571 |
| MERS | S2 | MH002341 | | Evolutioanry origin | Chiroptera | TGERKYR\|SA | 0.571 |
| MERS | S2 | MH002342 | | Evolutioanry origin | Chiroptera | TGERKYR\|SA | 0.571 |
| MERS | S2 | EF065509 | | Evolutioanry origin | Chiroptera | TGERKYR\|ST | 0.507 |
| MERS | S2 | EF065510 | | Evolutioanry origin | Chiroptera | TGERKYR\|ST | 0.507 |
| MERS | S2 | EF065511 | | Evolutioanry origin | Chiroptera | TGERKYR\|ST | 0.507 |
| MERS | S2 | EF065512 | | Evolutioanry origin | Chiroptera | TGERKYR\|ST | 0.507 |
| MERS | S2 | NC009020 | | Evolutioanry origin | Chiroptera | TGERKYR\|ST | 0.507 |
| MERS | S2 | KJ477103 | | Intermediate host | Artiodactyla | TGSRSAR\|SA | 0.707 |
| MERS | S2 | KU740200 | | Intermediate host | Artiodactyla | TGSRSAR\|SA | 0.707 |
| MERS | S2 | KY581695 | | Intermediate host | Artiodactyla | TGSRSAR\|SA | 0.707 |
| MERS | S2 | KY581696 | | Intermediate host | Artiodactyla | TGSRSAR\|SA | 0.707 |
| MERS | S2 | KY581697 | | Intermediate host | Artiodactyla | TGSRSAR\|SA | 0.707 |

| | | | | | | | |
|---|---|---|---|---|---|---|---|
| MERS | S2 | KY581698 | | Intermediate host | Artiodactyla | TGSRSAR\|SA | 0.707 |
| MERS | S2 | KY581699 | | Intermediate host | Artiodactyla | TGSRSAR\|SA | 0.707 |
| MERS | S2 | KY581700 | | Intermediate host | Artiodactyla | TGSRSAR\|SA | 0.707 |
| MERS | S2 | KY673149 | | Intermediate host | Artiodactyla | TGSRSAR\|SA | 0.707 |
| MERS | S2 | MF598594 | | Intermediate host | Artiodactyla | TGSRSAR\|SA | 0.707 |
| MERS | S2 | MF598595 | | Intermediate host | Artiodactyla | TGSRSAR\|SA | 0.707 |
| MERS | S2 | MF598596 | | Intermediate host | Artiodactyla | TGSRSAR\|SA | 0.707 |
| MERS | S2 | MF598597 | | Intermediate host | Artiodactyla | TGSRSAR\|SA | 0.707 |
| MERS | S2 | MF598598 | | Intermediate host | Artiodactyla | TGSRSAR\|SA | 0.707 |
| MERS | S2 | MF598599 | | Intermediate host | Artiodactyla | TGSRSAR\|SA | 0.707 |
| MERS | S2 | MF598600 | | Intermediate host | Artiodactyla | TGSRSAR\|SA | 0.707 |
| MERS | S2 | MF598601 | | Intermediate host | Artiodactyla | TGSRSAR\|SA | 0.707 |
| MERS | S2 | MF598602 | | Intermediate host | Artiodactyla | TGSRSAR\|SA | 0.707 |
| MERS | S2 | MF598603 | | Intermediate host | Artiodactyla | TGSRSAR\|SA | 0.707 |
| MERS | S2 | MF598604 | | Intermediate host | Artiodactyla | TGSRSAR\|SA | 0.707 |
| MERS | S2 | MF598605 | | Intermediate host | Artiodactyla | TGSRSAR\|SA | 0.707 |
| MERS | S2 | MF598606 | | Intermediate host | Artiodactyla | TGSRSAR\|SA | 0.707 |
| MERS | S2 | MF598607 | | Intermediate host | Artiodactyla | TGSRSAR\|SA | 0.707 |
| MERS | S2 | MF598608 | | Intermediate host | Artiodactyla | TGSRSAR\|SA | 0.707 |
| MERS | S2 | MF598609 | | Intermediate host | Artiodactyla | TGSRSAR\|SA | 0.707 |
| MERS | S2 | MF598610 | | Intermediate host | Artiodactyla | TGSRSAR\|SA | 0.707 |
| MERS | S2 | MF598611 | | Intermediate host | Artiodactyla | TGSRSAR\|SA | 0.707 |
| MERS | S2 | MF598612 | | Intermediate host | Artiodactyla | TGSRSAR\|SA | 0.707 |
| MERS | S2 | MF598613 | | Intermediate host | Artiodactyla | TGSRSAR\|SA | 0.707 |
| MERS | S2 | MF598614 | | Intermediate host | Artiodactyla | TGSRSAR\|SA | 0.707 |
| MERS | S2 | MF598615 | | Intermediate host | Artiodactyla | TGSRSAR\|SA | 0.707 |
| MERS | S2 | MF598616 | | Intermediate host | Artiodactyla | TGSRSAR\|SA | 0.707 |
| MERS | S2 | MF598617 | | Intermediate host | Artiodactyla | TGSRSAR\|SA | 0.707 |
| MERS | S2 | MF598618 | | Intermediate host | Artiodactyla | TGSRSAR\|SA | 0.707 |
| MERS | S2 | MF598619 | | Intermediate host | Artiodactyla | TGSRSAR\|SA | 0.707 |
| MERS | S2 | MF598620 | | Intermediate host | Artiodactyla | TGSRSAR\|SA | 0.707 |
| MERS | S2 | MF598621 | | Intermediate host | Artiodactyla | TGSRSAR\|SA | 0.707 |
| MERS | S2 | MF598622 | | Intermediate host | Artiodactyla | TGSRSAR\|SA | 0.707 |
| MERS | S2 | MF598623 | | Intermediate host | Artiodactyla | TGSRSAR\|SA | 0.707 |
| MERS | S2 | MF598624 | | Intermediate host | Artiodactyla | TGSRSAR\|SA | 0.707 |
| MERS | S2 | MF598625 | | Intermediate host | Artiodactyla | TGSRSAR\|SA | 0.707 |
| MERS | S2 | MF598626 | | Intermediate host | Artiodactyla | TGSRSAR\|SA | 0.707 |
| MERS | S2 | MF598627 | | Intermediate host | Artiodactyla | TGSRSAR\|SA | 0.707 |
| MERS | S2 | MF598628 | | Intermediate host | Artiodactyla | TGSRSAR\|SA | 0.707 |
| MERS | S2 | MF598629 | | Intermediate host | Artiodactyla | TGSRSAR\|SA | 0.707 |
| MERS | S2 | MF598630 | | Intermediate host | Artiodactyla | TGSRSAR\|SA | 0.707 |
| MERS | S2 | MF598631 | | Intermediate host | Artiodactyla | TGSRSAR\|SA | 0.707 |
| MERS | S2 | MF598632 | | Intermediate host | Artiodactyla | TGSRSAR\|SA | 0.707 |
| MERS | S2 | MF598633 | | Intermediate host | Artiodactyla | TGSRSAR\|SA | 0.707 |
| MERS | S2 | MF598634 | | Intermediate host | Artiodactyla | TGSRSAR\|SA | 0.707 |
| MERS | S2 | MF598635 | | Intermediate host | Artiodactyla | TGSRSAR\|SA | 0.707 |
| MERS | S2 | MF598636 | | Intermediate host | Artiodactyla | TGSRSAR\|SA | 0.707 |
| MERS | S2 | MF598637 | | Intermediate host | Artiodactyla | TGSRSAR\|SA | 0.707 |
| MERS | S2 | MF598638 | | Intermediate host | Artiodactyla | TGSRSAR\|SA | 0.707 |
| MERS | S2 | MF598639 | | Intermediate host | Artiodactyla | TGSRSAR\|SA | 0.707 |
| MERS | S2 | MF598640 | | Intermediate host | Artiodactyla | TGSRSAR\|SA | 0.707 |
| MERS | S2 | MF598641 | | Intermediate host | Artiodactyla | TGSRSAR\|SA | 0.707 |
| MERS | S2 | MF598642 | | Intermediate host | Artiodactyla | TGSRSAR\|SA | 0.707 |
| MERS | S2 | MF598643 | | Intermediate host | Artiodactyla | TGSRSAR\|SA | 0.707 |
| MERS | S2 | MF598644 | | Intermediate host | Artiodactyla | TGSRSAR\|SA | 0.707 |
| MERS | S2 | MF598645 | | Intermediate host | Artiodactyla | TGSRSAR\|SA | 0.707 |
| MERS | S2 | MF598646 | | Intermediate host | Artiodactyla | TGSRSAR\|SA | 0.707 |
| MERS | S2 | MF598647 | | Intermediate host | Artiodactyla | TGSRSAR\|SA | 0.707 |
| MERS | S2 | MF598648 | | Intermediate host | Artiodactyla | TGSRSAR\|SA | 0.707 |
| MERS | S2 | MF598649 | | Intermediate host | Artiodactyla | TGSRSAR\|SA | 0.707 |
| MERS | S2 | MF598650 | | Intermediate host | Artiodactyla | TGSRSAR\|SA | 0.707 |
| MERS | S2 | MF598651 | | Intermediate host | Artiodactyla | TGSRSAR\|SA | 0.707 |
| MERS | S2 | MF598652 | | Intermediate host | Artiodactyla | TGSRSAR\|SA | 0.707 |
| MERS | S2 | MF598653 | | Intermediate host | Artiodactyla | TGSRSAR\|SA | 0.707 |
| MERS | S2 | MF598654 | | Intermediate host | Artiodactyla | TGSRSAR\|SA | 0.707 |
| MERS | S2 | MF598655 | | Intermediate host | Artiodactyla | TGSRSAR\|SA | 0.707 |
| MERS | S2 | MF598656 | | Intermediate host | Artiodactyla | TGSRSAR\|SA | 0.707 |
| MERS | S2 | MF598657 | | Intermediate host | Artiodactyla | TGSRSAR\|SA | 0.707 |
| MERS | S2 | MF598658 | | Intermediate host | Artiodactyla | TGSRSAR\|SA | 0.707 |
| MERS | S2 | MF598659 | | Intermediate host | Artiodactyla | TGSRSAR\|SA | 0.707 |
| MERS | S2 | MF598660 | | Intermediate host | Artiodactyla | TGSRSAR\|SA | 0.707 |
| MERS | S2 | MF598661 | | Intermediate host | Artiodactyla | TGSRSAR\|SA | 0.707 |
| MERS | S2 | MF598662 | | Intermediate host | Artiodactyla | TGSRSAR\|SA | 0.707 |

| | | | | | | | |
|---|---|---|---|---|---|---|---|
| MERS | S2 | MF598663 | | Intermediate host | Artiodactyla | TGSRSAR\|SA | 0.707 |
| MERS | S2 | MF598664 | | Intermediate host | Artiodactyla | TGSRSAR\|SA | 0.707 |
| MERS | S2 | MF598665 | | Intermediate host | Artiodactyla | TGSRSAR\|SA | 0.707 |
| MERS | S2 | MF598666 | | Intermediate host | Artiodactyla | TGSRSAR\|SA | 0.707 |
| MERS | S2 | MF598667 | | Intermediate host | Artiodactyla | TGSRSAR\|SA | 0.707 |
| MERS | S2 | MF598668 | | Intermediate host | Artiodactyla | TGSRSAR\|SA | 0.707 |
| MERS | S2 | MF598669 | | Intermediate host | Artiodactyla | TGSRSAR\|SA | 0.707 |
| MERS | S2 | MF598670 | | Intermediate host | Artiodactyla | TGSRSAR\|SA | 0.707 |
| MERS | S2 | MF598671 | | Intermediate host | Artiodactyla | TGSRSAR\|SA | 0.707 |
| MERS | S2 | MF598672 | | Intermediate host | Artiodactyla | TGSRSAR\|SA | 0.707 |
| MERS | S2 | MF598673 | | Intermediate host | Artiodactyla | TGSRSAR\|SA | 0.707 |
| MERS | S2 | MF598674 | | Intermediate host | Artiodactyla | TGSRSAR\|SA | 0.707 |
| MERS | S2 | MF598675 | | Intermediate host | Artiodactyla | TGSRSAR\|SA | 0.707 |
| MERS | S2 | MF598676 | | Intermediate host | Artiodactyla | TGSRSAR\|SA | 0.707 |
| MERS | S2 | MF598677 | | Intermediate host | Artiodactyla | TGSRSAR\|SA | 0.707 |
| MERS | S2 | MF598678 | | Intermediate host | Artiodactyla | TGSRSAR\|SA | 0.707 |
| MERS | S2 | MF598679 | | Intermediate host | Artiodactyla | TGSRSAR\|SA | 0.707 |
| MERS | S2 | MF598680 | | Intermediate host | Artiodactyla | TGSRSAR\|SA | 0.707 |
| MERS | S2 | MF598681 | | Intermediate host | Artiodactyla | TGSRSAR\|SA | 0.707 |
| MERS | S2 | MF598682 | | Intermediate host | Artiodactyla | TGSRSAR\|SA | 0.707 |
| MERS | S2 | MF598683 | | Intermediate host | Artiodactyla | TGSRSAR\|SA | 0.707 |
| MERS | S2 | MF598684 | | Intermediate host | Artiodactyla | TGSRSAR\|SA | 0.707 |
| MERS | S2 | MF598685 | | Intermediate host | Artiodactyla | TGSRSAR\|SA | 0.707 |
| MERS | S2 | MF598686 | | Intermediate host | Artiodactyla | TGSRSAR\|SA | 0.707 |
| MERS | S2 | MF598687 | | Intermediate host | Artiodactyla | TGSRSAR\|SA | 0.707 |
| MERS | S2 | MF598688 | | Intermediate host | Artiodactyla | TGSRSAR\|SA | 0.707 |
| MERS | S2 | MF598689 | | Intermediate host | Artiodactyla | TGSRSAR\|SA | 0.707 |
| MERS | S2 | MF598690 | | Intermediate host | Artiodactyla | TGSRSAR\|SA | 0.707 |
| MERS | S2 | MF598691 | | Intermediate host | Artiodactyla | TGSRSAR\|SA | 0.707 |
| MERS | S2 | MF598692 | | Intermediate host | Artiodactyla | TGSRSAR\|SA | 0.707 |
| MERS | S2 | MF598693 | | Intermediate host | Artiodactyla | TGSRSAR\|SA | 0.707 |
| MERS | S2 | MF598694 | | Intermediate host | Artiodactyla | TGSRSAR\|SA | 0.707 |
| MERS | S2 | MF598695 | | Intermediate host | Artiodactyla | TGSRSAR\|SA | 0.707 |
| MERS | S2 | MF598696 | | Intermediate host | Artiodactyla | TGSRSAR\|SA | 0.707 |
| MERS | S2 | MF598697 | | Intermediate host | Artiodactyla | TGSRSAR\|SA | 0.707 |
| MERS | S2 | MF598698 | | Intermediate host | Artiodactyla | TGSRSAR\|SA | 0.707 |
| MERS | S2 | MF598699 | | Intermediate host | Artiodactyla | TGSRSAR\|SA | 0.707 |
| MERS | S2 | MF598700 | | Intermediate host | Artiodactyla | TGSRSAR\|SA | 0.707 |
| MERS | S2 | MF598701 | | Intermediate host | Artiodactyla | TGSRSAR\|SA | 0.707 |
| MERS | S2 | MF598702 | | Intermediate host | Artiodactyla | TGSRSAR\|SA | 0.707 |
| MERS | S2 | MF598703 | | Intermediate host | Artiodactyla | TGSRSAR\|SA | 0.707 |
| MERS | S2 | MF598704 | | Intermediate host | Artiodactyla | TGSRSAR\|SA | 0.707 |
| MERS | S2 | MF598705 | | Intermediate host | Artiodactyla | TGSRSAR\|SA | 0.707 |
| MERS | S2 | MF598706 | | Intermediate host | Artiodactyla | TGSRSAR\|SA | 0.707 |
| MERS | S2 | MF598707 | | Intermediate host | Artiodactyla | TGSRSAR\|SA | 0.707 |
| MERS | S2 | MF598708 | | Intermediate host | Artiodactyla | TGSRSAR\|SA | 0.707 |
| MERS | S2 | MF598709 | | Intermediate host | Artiodactyla | TGSRSAR\|SA | 0.707 |
| MERS | S2 | MF598710 | | Intermediate host | Artiodactyla | TGSRSAR\|SA | 0.707 |
| MERS | S2 | MF598711 | | Intermediate host | Artiodactyla | TGSRSAR\|SA | 0.707 |
| MERS | S2 | MF598712 | | Intermediate host | Artiodactyla | TGSRSAR\|SA | 0.707 |
| MERS | S2 | MF598713 | | Intermediate host | Artiodactyla | TGSRSAR\|SA | 0.707 |
| MERS | S2 | MF598714 | | Intermediate host | Artiodactyla | TGSRSAR\|SA | 0.707 |
| MERS | S2 | MF598715 | | Intermediate host | Artiodactyla | TGSRSAR\|SA | 0.707 |
| MERS | S2 | MF598716 | | Intermediate host | Artiodactyla | TGSRSAR\|SA | 0.707 |
| MERS | S2 | MF598717 | | Intermediate host | Artiodactyla | TGSRSAR\|SA | 0.707 |
| MERS | S2 | MF598718 | | Intermediate host | Artiodactyla | TGSRSAR\|SA | 0.707 |
| MERS | S2 | MF598719 | | Intermediate host | Artiodactyla | TGSRSAR\|SA | 0.707 |
| MERS | S2 | MF598720 | | Intermediate host | Artiodactyla | TGSRSAR\|SA | 0.707 |
| MERS | S2 | MF598721 | | Intermediate host | Artiodactyla | TGSRSAR\|SA | 0.707 |
| MERS | S2 | MF598722 | | Intermediate host | Artiodactyla | TGSRSAR\|SA | 0.707 |
| MERS | S2 | MG923466 | | Intermediate host | Artiodactyla | TGSRSAR\|SA | 0.707 |
| MERS | S2 | MG923467 | | Intermediate host | Artiodactyla | TGSRSAR\|SA | 0.707 |
| MERS | S2 | MG923468 | | Intermediate host | Artiodactyla | TGSRSAR\|SA | 0.707 |
| MERS | S2 | MG923470 | | Intermediate host | Artiodactyla | TGSRSAR\|SA | 0.707 |
| MERS | S2 | MG923471 | | Intermediate host | Artiodactyla | TGSRSAR\|SA | 0.707 |
| MERS | S2 | MG923472 | | Intermediate host | Artiodactyla | TGSRSAR\|SA | 0.707 |
| MERS | S2 | MG923473 | | Intermediate host | Artiodactyla | TGSRSAR\|SA | 0.707 |
| MERS | S2 | MG923474 | | Intermediate host | Artiodactyla | TGSRSAR\|SA | 0.707 |
| MERS | S2 | MG923475 | | Intermediate host | Artiodactyla | TGSRSAR\|SA | 0.707 |
| MERS | S2 | MG923476 | | Intermediate host | Artiodactyla | TGSRSAR\|SA | 0.707 |
| MERS | S2 | MG923477 | | Intermediate host | Artiodactyla | TGSRSAR\|SA | 0.707 |
| MERS | S2 | MG923478 | | Intermediate host | Artiodactyla | TGSRSAR\|SA | 0.707 |
| MERS | S2 | MG923479 | | Intermediate host | Artiodactyla | TGSRSAR\|SA | 0.707 |

| | | | | | | | |
|---|---|---|---|---|---|---|---|
| MERS | S2 | MG923480 | | Intermediate host | Artiodactyla | TGSRSAR\|SA | 0.707 |
| MERS | S2 | MG923481 | | Intermediate host | Artiodactyla | TGSRSAR\|SA | 0.707 |
| MERS | S2 | MH259485 | | Intermediate host | Artiodactyla | TGSRSAR\|SA | 0.707 |
| MERS | S2 | MH259486 | | Intermediate host | Artiodactyla | TGSRSAR\|SA | 0.707 |
| MERS | S2 | MH734114 | | Intermediate host | Artiodactyla | TGSRSAR\|SA | 0.707 |
| MERS | S2 | MH734115 | | Intermediate host | Artiodactyla | TGSRSAR\|SA | 0.707 |
| MERS | S2 | MK357908 | | Intermediate host | Artiodactyla | TGSRSAR\|SA | 0.707 |
| MERS | S2 | MK357909 | | Intermediate host | Artiodactyla | TGSRSAR\|SA | 0.707 |
| MERS | S2 | MK564474 | | Intermediate host | Artiodactyla | TGSRSAR\|SA | 0.707 |
| MERS | S2 | MK564475 | | Intermediate host | Artiodactyla | TGSRSAR\|SA | 0.707 |
| MERS | S2 | MK967708 | | Intermediate host | Artiodactyla | TGSRSAR\|SA | 0.707 |
| MERS | S2 | MN654970 | | Intermediate host | Artiodactyla | TGSRSAR\|SA | 0.707 |
| MERS | S2 | MN654971 | | Intermediate host | Artiodactyla | TGSRSAR\|SA | 0.707 |
| MERS | S2 | MN654972 | | Intermediate host | Artiodactyla | TGSRSAR\|SA | 0.707 |
| MERS | S2 | MN654973 | | Intermediate host | Artiodactyla | TGSRSAR\|SA | 0.707 |
| MERS | S2 | MN654974 | | Intermediate host | Artiodactyla | TGSRSAR\|SA | 0.707 |
| MERS | S2 | MN654975 | | Intermediate host | Artiodactyla | TGSRSAR\|SA | 0.707 |
| MERS | S2 | MN654976 | | Intermediate host | Artiodactyla | TGSRSAR\|SA | 0.707 |
| MERS | S2 | MN654977 | | Intermediate host | Artiodactyla | TGSRSAR\|SA | 0.707 |
| MERS | S2 | MN654978 | | Intermediate host | Artiodactyla | TGSRSAR\|SA | 0.707 |
| MERS | S2 | MN654979 | | Intermediate host | Artiodactyla | TGSRSAR\|SA | 0.707 |
| MERS | S2 | MN654980 | | Intermediate host | Artiodactyla | TGSRSAR\|SA | 0.707 |
| MERS | S2 | MN654981 | | Intermediate host | Artiodactyla | TGSRSAR\|SA | 0.707 |
| MERS | S2 | MN654982 | | Intermediate host | Artiodactyla | TGSRSAR\|SA | 0.707 |
| MERS | S2 | MN654983 | | Intermediate host | Artiodactyla | TGSRSAR\|SA | 0.707 |
| MERS | S2 | MN654984 | | Intermediate host | Artiodactyla | TGSRSAR\|SA | 0.707 |
| MERS | S2 | MN654985 | | Intermediate host | Artiodactyla | TGSRSAR\|SA | 0.707 |
| MERS | S2 | MN654986 | | Intermediate host | Artiodactyla | TGSRSAR\|SA | 0.707 |
| MERS | S2 | MN654987 | | Intermediate host | Artiodactyla | TGSRSAR\|SA | 0.707 |
| MERS | S2 | MN654988 | | Intermediate host | Artiodactyla | TGSRSAR\|SA | 0.707 |
| MERS | S2 | MN654989 | | Intermediate host | Artiodactyla | TGSRSAR\|SA | 0.707 |
| MERS | S2 | MN654990 | | Intermediate host | Artiodactyla | TGSRSAR\|SA | 0.707 |
| MERS | S2 | MN654991 | | Intermediate host | Artiodactyla | TGSRSAR\|SA | 0.707 |
| MERS | S2 | MN654992 | | Intermediate host | Artiodactyla | TGSRSAR\|SA | 0.707 |
| MERS | S2 | MN654993 | | Intermediate host | Artiodactyla | TGSRSAR\|SA | 0.707 |
| MERS | S2 | MN654994 | | Intermediate host | Artiodactyla | TGSRSAR\|SA | 0.707 |
| MERS | S2 | MN654995 | | Intermediate host | Artiodactyla | TGSRSAR\|SA | 0.707 |
| MERS | S2 | MN654996 | | Intermediate host | Artiodactyla | TGSRSAR\|SA | 0.707 |
| MERS | S2 | MN654997 | | Intermediate host | Artiodactyla | TGSRSAR\|SA | 0.707 |
| MERS | S2 | MN654998 | | Intermediate host | Artiodactyla | TGSRSAR\|SA | 0.707 |
| MERS | S2 | MN654999 | | Intermediate host | Artiodactyla | TGSRSAR\|SA | 0.707 |
| MERS | S2 | MN655000 | | Intermediate host | Artiodactyla | TGSRSAR\|SA | 0.707 |
| MERS | S2 | MN655001 | | Intermediate host | Artiodactyla | TGSRSAR\|SA | 0.707 |
| MERS | S2 | MN655002 | | Intermediate host | Artiodactyla | TGSRSAR\|SA | 0.707 |
| MERS | S2 | MN655003 | | Intermediate host | Artiodactyla | TGSRSAR\|SA | 0.707 |
| MERS | S2 | MN655004 | | Intermediate host | Artiodactyla | TGSRSAR\|SA | 0.707 |
| MERS | S2 | MN655005 | | Intermediate host | Artiodactyla | TGSRSAR\|SA | 0.707 |
| MERS | S2 | MN655006 | | Intermediate host | Artiodactyla | TGSRSAR\|SA | 0.707 |
| MERS | S2 | MN655007 | | Intermediate host | Artiodactyla | TGSRSAR\|SA | 0.707 |
| MERS | S2 | MN655008 | | Intermediate host | Artiodactyla | TGSRSAR\|SA | 0.707 |
| MERS | S2 | MN655009 | | Intermediate host | Artiodactyla | TGSRSAR\|SA | 0.707 |
| MERS | S2 | MN655010 | | Intermediate host | Artiodactyla | TGSRSAR\|SA | 0.707 |
| MERS | S2 | MN655011 | | Intermediate host | Artiodactyla | TGSRSAR\|SA | 0.707 |
| MERS | S2 | MN655012 | | Intermediate host | Artiodactyla | TGSRSAR\|SA | 0.707 |
| MERS | S2 | MN655013 | | Intermediate host | Artiodactyla | TGSRSAR\|SA | 0.707 |
| MERS | S2 | MN655014 | | Intermediate host | Artiodactyla | TGSRSAR\|SA | 0.707 |
| MERS | S2 | MN655015 | | Intermediate host | Artiodactyla | TGSRSAR\|SA | 0.707 |
| MERS | S2 | MN655016 | | Intermediate host | Artiodactyla | TGSRSAR\|SA | 0.707 |
| MERS | S2 | MN655017 | | Intermediate host | Artiodactyla | TGSRSAR\|SA | 0.707 |
| MERS | S2 | MN758606 | | Intermediate host | Artiodactyla | TGSRSAR\|SA | 0.707 |
| MERS | S2 | MN758607 | | Intermediate host | Artiodactyla | TGSRSAR\|SA | 0.707 |
| MERS | S2 | MN758608 | | Intermediate host | Artiodactyla | TGSRSAR\|SA | 0.707 |
| MERS | S2 | MN758609 | | Intermediate host | Artiodactyla | TGSRSAR\|SA | 0.707 |
| MERS | S2 | MN758610 | | Intermediate host | Artiodactyla | TGSRSAR\|SA | 0.707 |
| MERS | S2 | MN758611 | | Intermediate host | Artiodactyla | TGSRSAR\|SA | 0.707 |
| MERS | S2 | MN758612 | | Intermediate host | Artiodactyla | TGSRSAR\|SA | 0.707 |
| MERS | S2 | MN758613 | | Intermediate host | Artiodactyla | TGSRSAR\|SA | 0.707 |
| MERS | S2 | MN758614 | | Intermediate host | Artiodactyla | TGSRSAR\|SA | 0.707 |
| MERS | S2 | MT226600 | | Intermediate host | Artiodactyla | TGSRSAR\|SA | 0.707 |
| MERS | S2 | MT226601 | | Intermediate host | Artiodactyla | TGSRSAR\|SA | 0.707 |
| MERS | S2 | MT226602 | | Intermediate host | Artiodactyla | TGSRSAR\|SA | 0.707 |
| MERS | S2 | MT226603 | | Intermediate host | Artiodactyla | TGSRSAR\|SA | 0.707 |
| MERS | S2 | MT226604 | | Intermediate host | Artiodactyla | TGSRSAR\|SA | 0.707 |

| | | | | | | | |
|---|---|---|---|---|---|---|---|
| MERS | S2 | MT226605 | | Intermediate host | Artiodactyla | TGSRSAR\|SA | 0.707 |
| MERS | S2 | MT226606 | | Intermediate host | Artiodactyla | TGSRSAR\|SA | 0.707 |
| MERS | S2 | MT226607 | | Intermediate host | Artiodactyla | TGSRSAR\|SA | 0.707 |
| MERS | S2 | MW086527 | | Intermediate host | Artiodactyla | TGSRSAR\|SA | 0.707 |
| MERS | S2 | MW086528 | | Intermediate host | Artiodactyla | TGSRSAR\|SA | 0.707 |
| MERS | S2 | MW086529 | | Intermediate host | Artiodactyla | TGSRSAR\|SA | 0.707 |
| MERS | S2 | MW086530 | | Intermediate host | Artiodactyla | TGSRSAR\|SA | 0.707 |
| MERS | S2 | MW086531 | | Intermediate host | Artiodactyla | TGSRSAR\|SA | 0.707 |
| MERS | S2 | MW086532 | | Intermediate host | Artiodactyla | TGSRSAR\|SA | 0.707 |
| MERS | S2 | MW086533 | | Intermediate host | Artiodactyla | TGSRSAR\|SA | 0.707 |
| MERS | S2 | MW086534 | | Intermediate host | Artiodactyla | TGSRSAR\|SA | 0.707 |
| MERS | S2 | MW086535 | | Intermediate host | Artiodactyla | TGSRSAR\|SA | 0.707 |
| MERS | S2 | MW086538 | | Intermediate host | Artiodactyla | TGSRSAR\|SA | 0.707 |
| MERS | S2 | MW086539 | | Intermediate host | Artiodactyla | TGSRSAR\|SA | 0.707 |
| MERS | S2 | MW545527 | | Intermediate host | Artiodactyla | TGSRSAR\|SA | 0.707 |
| MERS | S2 | MW545528 | | Intermediate host | Artiodactyla | TGSRSAR\|SA | 0.707 |
| MERS | S2 | MH371127 | MERS-CoV | | Primates | TGSRSAR\|SA | 0.707 |
| MERS | S2 | NC019843 | MERS-CoV | | Primates | TGSRSAR\|SA | 0.707 |
| MERS | S2 | NC038294 | MERS-CoV | | Primates | TGSRSAR\|SA | 0.707 |
| NL63r | S2 | NC005831 | HCoV-NL63 | | Primates | LPQRNIR\|SS | 0.519 |
| OC43 | S1S2 | KM349742 | | Evolutioanry origin | Rodentia | STWRAKR\|DL | 0.749 |
| OC43 | S1S2 | KM349743 | | Evolutioanry origin | Rodentia | STWRAKR\|DL | 0.749 |
| OC43 | S1S2 | KM349744 | | Evolutioanry origin | Rodentia | STWRAKR\|DL | 0.749 |
| OC43 | S1S2 | MT820628 | | Evolutioanry origin | Rodentia | ATRRAKR\|DL | 0.761 |
| OC43 | S1S2 | MT820629 | | Evolutioanry origin | Rodentia | ATRRAKR\|DL | 0.761 |
| OC43 | S1S2 | MT820630 | | Evolutioanry origin | Rodentia | ATRRSKR\|DL | 0.793 |
| OC43 | S1S2 | MT820631 | | Evolutioanry origin | Rodentia | ATRRAKR\|DL | 0.761 |
| OC43 | S1S2 | NC026011 | | Evolutioanry origin | Rodentia | STWRAKR\|DL | 0.749 |
| OC43 | S1S2 | AB354579 | | Intermediate host | Artiodactyla | TKRRSRR\|AI | 0.776 |
| OC43 | S1S2 | AF220295 | | Intermediate host | Artiodactyla | TKRRSRR\|AI | 0.776 |
| OC43 | S1S2 | AF391541 | | Intermediate host | Artiodactyla | TKRRSRR\|SI | 0.851 |
| OC43 | S1S2 | AF391542 | | Intermediate host | Artiodactyla | TKRRSRR\|SI | 0.851 |
| OC43 | S1S2 | DQ011855 | | Intermediate host | Artiodactyla | TALRSRR\|SF | 0.715 |
| OC43 | S1S2 | DQ811784 | | Intermediate host | Artiodactyla | TKRRSRR\|SI | 0.851 |
| OC43 | S1S2 | DQ915164 | | Intermediate host | Artiodactyla | TKRRSRR\|SI | 0.851 |
| OC43 | S1S2 | EF424615 | | Intermediate host | Artiodactyla | TKRRSRR\|SI | 0.851 |
| OC43 | S1S2 | EF424617 | | Intermediate host | Artiodactyla | TKRRSRR\|SI | 0.851 |
| OC43 | S1S2 | EF424619 | | Intermediate host | Artiodactyla | TKRRSRR\|SI | 0.851 |
| OC43 | S1S2 | EF424620 | | Intermediate host | Artiodactyla | TKRRSRR\|SI | 0.851 |
| OC43 | S1S2 | EF424621 | | Intermediate host | Artiodactyla | TKRRSRR\|SI | 0.851 |
| OC43 | S1S2 | EF424622 | | Intermediate host | Artiodactyla | TKRRSRR\|SI | 0.851 |
| OC43 | S1S2 | EF424623 | | Intermediate host | Artiodactyla | TKRRSRR\|SI | 0.851 |
| OC43 | S1S2 | EF424624 | | Intermediate host | Artiodactyla | TKRRSRR\|SI | 0.851 |
| OC43 | S1S2 | FJ425184 | | Intermediate host | Artiodactyla | TKRRSRR\|SI | 0.851 |
| OC43 | S1S2 | FJ425185 | | Intermediate host | Artiodactyla | TKRRSRR\|SI | 0.851 |
| OC43 | S1S2 | FJ425186 | | Intermediate host | Artiodactyla | TKRRSRR\|SI | 0.851 |
| OC43 | S1S2 | FJ425187 | | Intermediate host | Artiodactyla | TKRRSRR\|SI | 0.851 |
| OC43 | S1S2 | FJ425188 | | Intermediate host | Artiodactyla | TKRRSRR\|SI | 0.851 |
| OC43 | S1S2 | FJ425189 | | Intermediate host | Artiodactyla | TKRRSRR\|SI | 0.851 |
| OC43 | S1S2 | FJ938063 | | Intermediate host | Artiodactyla | TKRRSRR\|SI | 0.851 |
| OC43 | S1S2 | FJ938064 | | Intermediate host | Artiodactyla | TKRRSRR\|SI | 0.851 |
| OC43 | S1S2 | FJ938065 | | Intermediate host | Artiodactyla | TKRRSRR\|SI | 0.851 |
| OC43 | S1S2 | FJ938066 | | Intermediate host | Artiodactyla | TKRRSRR\|SI | 0.851 |
| OC43 | S1S2 | KF906249 | | Intermediate host | Artiodactyla | IDRRSRR\|AI | 0.719 |
| OC43 | S1S2 | KF906250 | | Intermediate host | Artiodactyla | IDRRSRR\|AI | 0.719 |
| OC43 | S1S2 | KU558922 | | Intermediate host | Artiodactyla | IKRRSRR\|SI | 0.829 |
| OC43 | S1S2 | KU558923 | | Intermediate host | Artiodactyla | IKRRSRR\|SI | 0.829 |
| OC43 | S1S2 | KU886219 | | Intermediate host | Artiodactyla | TKRRSRR\|SI | 0.851 |
| OC43 | S1S2 | KX982264 | | Intermediate host | Artiodactyla | TKRRSRR\|AI | 0.776 |
| OC43 | S1S2 | KY419103 | | Intermediate host | Artiodactyla | TALRSRR\|SF | 0.715 |
| OC43 | S1S2 | KY419104 | | Intermediate host | Artiodactyla | TALRSRR\|SF | 0.715 |
| OC43 | S1S2 | KY419105 | | Intermediate host | Artiodactyla | TALRSRR\|SF | 0.715 |
| OC43 | S1S2 | KY419106 | | Intermediate host | Artiodactyla | TSLRSRR\|SL | 0.759 |
| OC43 | S1S2 | KY419107 | | Intermediate host | Artiodactyla | TSLRSRR\|SL | 0.759 |
| OC43 | S1S2 | KY419109 | | Intermediate host | Artiodactyla | TSLRSRR\|SL | 0.759 |
| OC43 | S1S2 | KY419110 | | Intermediate host | Artiodactyla | TALRSRR\|SF | 0.715 |
| OC43 | S1S2 | KY419112 | | Intermediate host | Artiodactyla | TALRSRR\|SF | 0.715 |
| OC43 | S1S2 | KY419113 | | Intermediate host | Artiodactyla | TSLRSRR\|SL | 0.759 |
| OC43 | S1S2 | KY994645 | | Intermediate host | Artiodactyla | TALRSRR\|SF | 0.715 |
| OC43 | S1S2 | LC494126 | | Intermediate host | Artiodactyla | TKRRSRR\|SI | 0.851 |
| OC43 | S1S2 | LC494127 | | Intermediate host | Artiodactyla | TKRRSRR\|SI | 0.851 |
| OC43 | S1S2 | LC494128 | | Intermediate host | Artiodactyla | TKRRSRR\|SI | 0.851 |
| OC43 | S1S2 | LC494129 | | Intermediate host | Artiodactyla | TKRRSRR\|SI | 0.851 |

| | | | | | | | |
|---|---|---|---|---|---|---|---|
| OC43 | S1S2 | LC494130 | | Intermediate host | Artiodactyla | TKRRSRR\|SI | 0.851 |
| OC43 | S1S2 | LC494131 | | Intermediate host | Artiodactyla | TKRRSRR\|SI | 0.851 |
| OC43 | S1S2 | LC494132 | | Intermediate host | Artiodactyla | TKRRSRR\|SI | 0.851 |
| OC43 | S1S2 | LC494133 | | Intermediate host | Artiodactyla | TKRRSRR\|SI | 0.851 |
| OC43 | S1S2 | LC494134 | | Intermediate host | Artiodactyla | TKRRSRR\|SI | 0.851 |
| OC43 | S1S2 | LC494135 | | Intermediate host | Artiodactyla | TKRRSRR\|SI | 0.851 |
| OC43 | S1S2 | LC494136 | | Intermediate host | Artiodactyla | TKRRSRR\|SI | 0.851 |
| OC43 | S1S2 | LC494137 | | Intermediate host | Artiodactyla | TKRRSRR\|SI | 0.851 |
| OC43 | S1S2 | LC494138 | | Intermediate host | Artiodactyla | TKRRSRR\|SI | 0.851 |
| OC43 | S1S2 | LC494139 | | Intermediate host | Artiodactyla | TKRRSRR\|SI | 0.851 |
| OC43 | S1S2 | LC494140 | | Intermediate host | Artiodactyla | TKRRSRR\|SI | 0.851 |
| OC43 | S1S2 | LC494141 | | Intermediate host | Artiodactyla | TKRRSRR\|SI | 0.851 |
| OC43 | S1S2 | LC494142 | | Intermediate host | Artiodactyla | TKRRSRR\|SI | 0.851 |
| OC43 | S1S2 | LC494143 | | Intermediate host | Artiodactyla | TKRRSRR\|SI | 0.851 |
| OC43 | S1S2 | LC494144 | | Intermediate host | Artiodactyla | TKRRSRR\|SI | 0.851 |
| OC43 | S1S2 | LC494145 | | Intermediate host | Artiodactyla | TKRRSRR\|SI | 0.851 |
| OC43 | S1S2 | LC494147 | | Intermediate host | Artiodactyla | TKRRSRR\|SI | 0.851 |
| OC43 | S1S2 | LC494148 | | Intermediate host | Artiodactyla | TKRRSRR\|SI | 0.851 |
| OC43 | S1S2 | LC494149 | | Intermediate host | Artiodactyla | TKRRSRR\|SI | 0.851 |
| OC43 | S1S2 | LC494150 | | Intermediate host | Artiodactyla | TKRRSRR\|SI | 0.851 |
| OC43 | S1S2 | LC494151 | | Intermediate host | Artiodactyla | TKRRSRR\|SI | 0.851 |
| OC43 | S1S2 | LC494152 | | Intermediate host | Artiodactyla | TKRRSRR\|SI | 0.851 |
| OC43 | S1S2 | LC494153 | | Intermediate host | Artiodactyla | TKRRSRR\|SI | 0.851 |
| OC43 | S1S2 | LC494154 | | Intermediate host | Artiodactyla | TKRRSRR\|SI | 0.851 |
| OC43 | S1S2 | LC494155 | | Intermediate host | Artiodactyla | TKRRSRR\|SI | 0.851 |
| OC43 | S1S2 | LC494156 | | Intermediate host | Artiodactyla | TKRRSRR\|SI | 0.851 |
| OC43 | S1S2 | LC494157 | | Intermediate host | Artiodactyla | TKRRSRR\|SI | 0.851 |
| OC43 | S1S2 | LC494158 | | Intermediate host | Artiodactyla | TKRRSRR\|SI | 0.851 |
| OC43 | S1S2 | LC494159 | | Intermediate host | Artiodactyla | TKRRSRR\|SI | 0.851 |
| OC43 | S1S2 | LC494160 | | Intermediate host | Artiodactyla | TKRRSRR\|SI | 0.851 |
| OC43 | S1S2 | LC494161 | | Intermediate host | Artiodactyla | TKRRSRR\|SI | 0.851 |
| OC43 | S1S2 | LC494162 | | Intermediate host | Artiodactyla | TKRRSRR\|SI | 0.851 |
| OC43 | S1S2 | LC494163 | | Intermediate host | Artiodactyla | TKRRSRR\|SI | 0.851 |
| OC43 | S1S2 | LC494164 | | Intermediate host | Artiodactyla | TKRRSRR\|SI | 0.851 |
| OC43 | S1S2 | LC494165 | | Intermediate host | Artiodactyla | TKRRSRR\|SI | 0.851 |
| OC43 | S1S2 | LC494166 | | Intermediate host | Artiodactyla | TKRRSRR\|SI | 0.851 |
| OC43 | S1S2 | LC494167 | | Intermediate host | Artiodactyla | TKRRSRR\|SI | 0.851 |
| OC43 | S1S2 | LC494168 | | Intermediate host | Artiodactyla | TKRRSRR\|SI | 0.851 |
| OC43 | S1S2 | LC494169 | | Intermediate host | Artiodactyla | TKRRSRR\|SI | 0.851 |
| OC43 | S1S2 | LC494170 | | Intermediate host | Artiodactyla | TKRRSRR\|SI | 0.851 |
| OC43 | S1S2 | LC494171 | | Intermediate host | Artiodactyla | TKRRSRR\|SI | 0.851 |
| OC43 | S1S2 | LC494172 | | Intermediate host | Artiodactyla | TKRRSRR\|SI | 0.851 |
| OC43 | S1S2 | LC494173 | | Intermediate host | Artiodactyla | TKRRSRR\|SI | 0.851 |
| OC43 | S1S2 | LC494174 | | Intermediate host | Artiodactyla | TKRRSRR\|SI | 0.851 |
| OC43 | S1S2 | LC494175 | | Intermediate host | Artiodactyla | TKRRSRR\|SI | 0.851 |
| OC43 | S1S2 | LC494176 | | Intermediate host | Artiodactyla | TKRRSRR\|SI | 0.851 |
| OC43 | S1S2 | LC494177 | | Intermediate host | Artiodactyla | TKRRSRR\|SI | 0.851 |
| OC43 | S1S2 | LC494178 | | Intermediate host | Artiodactyla | TKRRSRR\|SI | 0.851 |
| OC43 | S1S2 | LC494179 | | Intermediate host | Artiodactyla | TKRRSRR\|SI | 0.851 |
| OC43 | S1S2 | LC494180 | | Intermediate host | Artiodactyla | TKRRSRR\|SI | 0.851 |
| OC43 | S1S2 | LC494181 | | Intermediate host | Artiodactyla | TKRRSRR\|SI | 0.851 |
| OC43 | S1S2 | LC494182 | | Intermediate host | Artiodactyla | TKRRSRR\|SI | 0.851 |
| OC43 | S1S2 | LC494183 | | Intermediate host | Artiodactyla | TKRRSRR\|SI | 0.851 |
| OC43 | S1S2 | LC494184 | | Intermediate host | Artiodactyla | TKRRSRR\|SI | 0.851 |
| OC43 | S1S2 | LC494185 | | Intermediate host | Artiodactyla | TKRRSRR\|SI | 0.851 |
| OC43 | S1S2 | LC494186 | | Intermediate host | Artiodactyla | TKRRSRR\|SI | 0.851 |
| OC43 | S1S2 | LC494187 | | Intermediate host | Artiodactyla | TKRRSRR\|SI | 0.851 |
| OC43 | S1S2 | LC494188 | | Intermediate host | Artiodactyla | TKRRSRR\|SI | 0.851 |
| OC43 | S1S2 | LC494189 | | Intermediate host | Artiodactyla | TKRRSRR\|SI | 0.851 |
| OC43 | S1S2 | LC494190 | | Intermediate host | Artiodactyla | TKRRSRR\|SI | 0.851 |
| OC43 | S1S2 | LC494191 | | Intermediate host | Artiodactyla | TKRRSRR\|SI | 0.851 |
| OC43 | S1S2 | LC494192 | | Intermediate host | Artiodactyla | TKRRSRR\|SI | 0.851 |
| OC43 | S1S2 | MF083115 | | Intermediate host | Artiodactyla | TALRSRR\|SF | 0.715 |
| OC43 | S1S2 | MG518518 | | Intermediate host | Artiodactyla | TKRRSRR\|SI | 0.851 |
| OC43 | S1S2 | MG757138 | | Intermediate host | Artiodactyla | TKRRSRR\|AI | 0.776 |
| OC43 | S1S2 | MG757139 | | Intermediate host | Artiodactyla | TKRRSRR\|AI | 0.776 |
| OC43 | S1S2 | MG757140 | | Intermediate host | Artiodactyla | TKRRSRR\|AI | 0.776 |
| OC43 | S1S2 | MG757141 | | Intermediate host | Artiodactyla | TKRRSRR\|AI | 0.776 |
| OC43 | S1S2 | MG757142 | | Intermediate host | Artiodactyla | TKRRSRR\|AI | 0.776 |
| OC43 | S1S2 | MH043952 | | Intermediate host | Artiodactyla | TKRRSRR\|SI | 0.851 |
| OC43 | S1S2 | MH043953 | | Intermediate host | Artiodactyla | TKRRSRR\|SI | 0.851 |
| OC43 | S1S2 | MH043954 | | Intermediate host | Artiodactyla | TKRRSRR\|SI | 0.851 |
| OC43 | S1S2 | MH043955 | | Intermediate host | Artiodactyla | TKRRSRR\|SI | 0.851 |

| | | | | | | | |
|---|---|---|---|---|---|---|---|
| OC43 | S1S2 | MH810163 | | Intermediate host | Artiodactyla | TKRRSRR\|AI | 0.813 |
| OC43 | S1S2 | MN514962 | | Intermediate host | Artiodactyla | KERRSRR\|AI | 0.728 |
| OC43 | S1S2 | MN514963 | | Intermediate host | Artiodactyla | TDRRSRR\|AV | 0.812 |
| OC43 | S1S2 | MN514964 | | Intermediate host | Artiodactyla | TDRRSRR\|AI | 0.752 |
| OC43 | S1S2 | MN514965 | | Intermediate host | Artiodactyla | TDRRXRR\|AI | 0.738 |
| OC43 | S1S2 | MN514966 | | Intermediate host | Artiodactyla | TDRRSRR\|AV | 0.814 |
| OC43 | S1S2 | MN514967 | | Intermediate host | Artiodactyla | TDRRSRR\|AI | 0.752 |
| OC43 | S1S2 | MN982198 | | Intermediate host | Artiodactyla | TKRRSRR\|SI | 0.851 |
| OC43 | S1S2 | MN982199 | | Intermediate host | Artiodactyla | TKRRSRR\|SI | 0.851 |
| OC43 | S1S2 | MW165134 | | Intermediate host | Artiodactyla | TALRSRR\|SF | 0.715 |
| OC43 | S1S2 | MW711287 | | Intermediate host | Artiodactyla | TKRRSRR\|SI | 0.851 |
| OC43 | S1S2 | NC003045 | | Intermediate host | Artiodactyla | TKRRSRR\|SI | 0.851 |
| OC43 | S1S2 | U00735 | | Intermediate host | Artiodactyla | TKRRSRR\|AI | 0.776 |
| OC43 | S1S2 | JX860640 | | Intermediate host | Carnivora | TKRRSRR\|SI | 0.851 |
| OC43 | S1S2 | KX432213 | | Intermediate host | Carnivora | TKRRSRR\|SI | 0.851 |
| OC43 | S1S2 | NC017083 | | Intermediate host | Lagomorpha | TQLRSRR\|AI | 0.627 |
| OC43 | S1S2 | EF446615 | | Intermediate host | Perissodactyl a | TARRQRR\|SP | 0.819 |
| OC43 | S1S2 | LC061272 | | Intermediate host | Perissodactyl a | IARRQRR\|SP | 0.795 |
| OC43 | S1S2 | LC061273 | | Intermediate host | Perissodactyl a | TARRQRR\|SP | 0.819 |
| OC43 | S1S2 | LC061274 | | Intermediate host | Perissodactyl a | TARRQRR\|SP | 0.819 |
| OC43 | S1S2 | LC592689 | | Intermediate host | Perissodactyl a | TARRQRR\|SP | 0.819 |
| OC43 | S1S2 | FJ415324 | Human enteric CoV 4408 | | Primates | TKRRSRR\|AI | 0.776 |
| OC43 | S1S2 | FJ938067 | Human enteric CoV 4408 | | Primates | TKRRSRR\|AI | 0.776 |
| OC43 | S1S2 | MH685718 | HCoV-OC43 | | Primates | KNRRSRR\|AI | 0.751 |
| OC43 | S1S2 | NC006213 | HCoV-OC43 | | Primates | SKNRRSR\|GA | 0.529 |
| SARS-related | S1S2 | MT019529 | SARS-CoV-2 | | Primates | NSPRRAR\|SV | 0.620 |

Accession numbers of viruses tested in ProP:

<u>HCoV-229E:</u> 41 Sequences (Hosts: 1 Primate, 6 Chiroptera, 34 Artiodactyla); JQ410000, KT253269, KT253270, KT253271, KT253272, KT253324, KT253325, KT253326, KT253327, KT253328, KT368892, KT368893, KT368894, KT368895, KT368896, KT368897, KT368898, KT368899, KT368900, KT368901, KT368902, KT368903, KT368904, KT368905, KT368906, KT368907, KT368908, KT368909, KT368910, KT368911, KT368912, KT368913, KT368914, KT368915, KT368916, KU291449, KY073747, KY073748, MF593473, NC002645, NC028752

<u>MERS-CoV:</u> 276 Sequences (Hosts: 3 Primates, 241 Artiodactyla, 29 Chiroptera, 3 Eulipotyphla); DQ648794, EF065505, EF065506, EF065507, EF065508, EF065509, EF065510, EF065511, EF065512, KC545386, KC869678, KJ473820, KJ473821, KJ473822, KJ477103, KU740200, KY581695, KY581696, KY581697, KY581698, KY581699, KY581700, KY673149, MF593268, MF598594, MF598595, MF598596, MF598597, MF598598, MF598599, MF598600, MF598601, MF598602, MF598603, MF598604, MF598605, MF598606, MF598607, MF598608, MF598609, MF598610, MF598611, MF598612, MF598613, MF598614, MF598615, MF598616, MF598617,

MF598618, MF598619, MF598620, MF598621, MF598622, MF598623, MF598624, MF598625, MF598626, MF598627, MF598628, MF598629, MF598630, MF598631, MF598632, MF598633, MF598634, MF598635, MF598636, MF598637, MF598638, MF598639, MF598640, MF598641, MF598642, MF598643, MF598644, MF598645, MF598646, MF598647, MF598648, MF598649, MF598650, MF598651, MF598652, MF598653, MF598654, MF598655, MF598656, MF598657, MF598658, MF598659, MF598660, MF598661, MF598662, MF598663, MF598664, MF598665, MF598666, MF598667, MF598668, MF598669, MF598670, MF598671, MF598672, MF598673, MF598674, MF598675, MF598676, MF598677, MF598678, MF598679, MF598680, MF598681, MF598682, MF598683, MF598684, MF598685, MF598686, MF598687, MF598688, MF598689, MF598690, MF598691, MF598692, MF598693, MF598694, MF598695, MF598696, MF598697, MF598698, MF598699, MF598700, MF598701, MF598702, MF598703, MF598704, MF598705, MF598706, MF598707, MF598708, MF598709, MF598710, MF598711, MF598712, MF598713, MF598714, MF598715, MF598716, MF598717, MF598718, MF598719, MF598720, MF598721, MF598722, MG021451, MG021452, MG596802, MG596803, MG923466, MG923467, MG923468, MG923469, MG923470, MG923471, MG923472, MG923473, MG923474, MG923475, MG923476, MG923477, MG923478, MG923479, MG923480, MG923481, MG987420, MG987421, MH002337, MH002338, MH002339, MH002340, MH002341, MH002342, MH259485, MH259486, MH371127, MH734114, MH734115, MK357908, MK357909, MK564474, MK564475, MK679660, MK967708, MN654970, MN654971, MN654972, MN654973, MN654974, MN654975, MN654976, MN654977, MN654978, MN654979, MN654980, MN654981, MN654982, MN654983, MN654984, MN654985, MN654986, MN654987, MN654988, MN654989, MN654990, MN654991, MN654992, MN654993, MN654994, MN654995, MN654996, MN654997, MN654998, MN654999, MN655000, MN655001, MN655002, MN655003, MN655004, MN655005, MN655006, MN655007, MN655008, MN655009, MN655010, MN655011, MN655012, MN655013, MN655014, MN655015, MN655016, MN655017, MN758606, MN758607, MN758608, MN758609, MN758610, MN758611, MN758612, MN758613, MN758614, MT226600, MT226601, MT226602, MT226603, MT226604, MT226605, MT226606, MT226607, MW086527, MW086528, MW086529, MW086530, MW086531, MW086532,

MW086533, MW086534, MW086535, MW086538, MW086539, MW218395, MW545527, MW545528, NC009019, NC009020, NC019843, NC038294, NC039207

HCoV-NL63: 6 Sequences (Hosts: 1 Primate, 5 Chiroptera); KY073744, KY073745, KY073746, NC32107, NC48216, NC5831

HCoV-OC43: 151 sequences (Hosts: 4 Primates, 131 Artiodactyla, 8 Rodentia, 5 Perissodactyla, 2 Carnivora, 1 Lagomorpha); AB354579, AF220295, AF391541, AF391542, DQ011855, DQ811784, DQ915164, EF424615, EF424617, EF424619, EF424620, EF424621, EF424622, EF424623, EF424624, EF446615, FJ415324, FJ425184, FJ425185, FJ425186, FJ425187, FJ425188, FJ425189, FJ938063, FJ938064, FJ938065, FJ938066, FJ938067, JX860640, KF906249, KF906250, KM349742, KM349743, KM349744, KU558922, KU558923, KU886219, KX432213, KX982264, KY419103, KY419104, KY419105, KY419106, KY419107, KY419109, KY419110, KY419112, KY419113, KY994645, LC061272, LC061273, LC061274, LC494126, LC494127, LC494128, LC494129, LC494130, LC494131, LC494132, LC494133, LC494134, LC494135, LC494136, LC494137, LC494138, LC494139, LC494140, LC494141, LC494142, LC494143, LC494144, LC494145, LC494147, LC494148, LC494149, LC494150, LC494151, LC494152, LC494153, LC494154, LC494155, LC494156, LC494157, LC494158, LC494159, LC494160, LC494161, LC494162, LC494163, LC494164, LC494165, LC494166, LC494167, LC494168, LC494169, LC494170, LC494171, LC494172, LC494173, LC494174, LC494175, LC494176, LC494177, LC494178, LC494179, LC494180, LC494181, LC494182, LC494183, LC494184, LC494185, LC494186, LC494187, LC494188, LC494189, LC494190, LC494191, LC494192, LC592689, MF083115, MG518518, MG757138, MG757139, MG757140, MG757141, MG757142, MH043952, MH043953, MH043954, MH043955, MH685718, MH810163, MN514962, MN514963, MN514964, MN514965, MN514966, MN514967, MN982198, MN982199, MT820628, MT820629, MT820630, MT820631, MW165134, MW711287, NC026011, NC0003045, NC006213, NC017083, U00735

HCoV-HKU1: 31 sequences (Hosts: 1 Primate, 30 Rodentia); AB551247, AC000192, AF201929, FJ647218, FJ647219, FJ647220, FJ647221, FJ647222, FJ647223, FJ647224, FJ647225, FJ647226, FJ647227, FJ884687, FJ938068, GU593319, JF792616, JF792617, JQ173883, JX169866, JX169867,

KF850449, MF416379, MF618252, MF618253, MW620427, MW773844, NC001846, NC006577, NC012936, NC048217

SARS-related CoV: 82 sequences (Hosts: 2 Primates, 62 Chiroptera, 12 Carnivora, 6 Pholidota); AY304486, AY304488, AY545914, AY545915, AY545916, AY545917, AY545918, AY545919, AY572034, AY572035, AY572038, DQ071615, DQ412042, DQ412043, DQ648856, DQ648857, FJ588686, FJ959407, GU190215, JX993987, JX993988, KC881005, KC881006, KF294457, KF367457, KF569996, KJ473811, KJ473812, KJ473813, KJ473814, KJ473815, KJ473816, KP886808, KP886809, KT444582, KU182964, KY352407, KY417142, KY417143, KY417144, KY417145, KY417146, KY417147, KY417148, KY417149, KY417150, KY417151, KY417152, KY770858, KY770859, KY770860, KY938558, LC556375, MG772933, MG772934, MK211374, MK211375, MK211376, MK211377, MK211378, MN996532, MT019529, MT040333, MT040334, MT040335, MT040336, MT072864, MT121216, MT726043, MT726045, MW251308, MW703458, MZ081376, MZ081377, MZ081378, MZ081379, MZ081380, MZ081381, MZ081382, MZ190137, MZ190138, NC004718
